# Metabolic reprogramming of the infant gut by bifidobacteria-based probiotics drives exclusion of antibiotic-resistant pathobionts

**DOI:** 10.1101/2025.08.15.670594

**Authors:** Ahmed Bargheet, Gaute Hovde Bø, Museveni Justine, Sabrina John Moyo, Iren Høyland Löhr, Bjørn Blomberg, Nina Langeland, Claus Klingenberg, Veronika Kuchařová Pettersen

## Abstract

Early-life probiotics have emerged as a promising strategy to combat the global emergency of antimicrobial resistance by enhancing gut resilience in infants. However, how exactly probiotic bacteria affect resistant opportunistic pathogens, *i.e*., pathobionts, remains nascent. We investigated effects of probiotic supplementation in 152 full-term, healthy Tanzanian infants, a sub-cohort of the ProRIDE trial (NCT04172012). Administration of oral probiotics during the first 4 weeks of life led to enhanced gut colonization by the probiotic *Bifidobacterium* species while suppressing pathobionts such as extended-spectrum β-lactamase-producing *Enterobacterales* (ESBL-E). Integration of metagenomics with metabolomics revealed that probiotics decreased resistome load and mobilome richness at 6 weeks, with concurrent shifts in metabolome. Specifically, the intervention increased fecal lactate and pyruvate and reduced cross-feeding pathways leading to propionate and butyrate, which partly explained the reduced ESBL-E carriage. Our findings provide mechanistic insights into how probiotic-driven *Bifidobacterium* colonization modulates infant gut ecosystem to attenuate antimicrobial resistance in infancy.

## INTRODUCTION

Antimicrobial resistance (AMR) is a growing and persistent threat to global health. As a prime example, systemic infections caused by extended-spectrum β-lactamase-producing *Enterobacterales* (ESBL-E), such as *Escherichia coli* and *Klebsiella pneumoniae*, undermine empirical antibiotic therapy and lead to high mortality rates in low- and middle-income countries (LMIC)^1,2^. In Tanzania, for instance, bloodstream infections with ESBL-E are associated with a case fatality rate as high as 70%, compared to around 30% for infections caused by susceptible strains^3,4^. This example underscores the need for preventive strategies against AMR infections. And even though not all ESBL-E and other drug-resistant bacteria colonizing the human gut will cause a life-threatening infection, their early establishment may act as a long-term reservoir of AMR that can cause harm at any time later in life^5^.

The gut microbiome represents the first line of defense against colonization by opportunistic pathogens, *i.e.,* pathobionts, including AMR bacteria^6^. As a result, microbial interventions, particularly probiotic supplementation, are increasingly explored as tools to reinforce colonization resistance during infancy^6,7^. *Bifidobacterium* species, abundant in breastfed infants, are key bacterial symbionts that modulate immune development and outcompete taxa with pathogenic potential^8^. Notably, supplementation with *Bifidobacterium longum* has been shown to reduce gut colonization by *Enterobacterales*^9^, major vectors for the carriage of AMR genes. Complementary findings from preterm infant cohorts suggest that probiotics may also reduce colonization with other AMR-associated taxa^10,11^. However, mechanistic insights remain limited.

Beyond taxonomy, the gut microbiome harbors a dynamic resistome, *i.e*., the repertoire of antibiotic resistance genes (ARGs), and a mobilome consisting of mobile genetic elements (MGEs), such as plasmids and transposons, that facilitate horizontal gene transfer^12^. These elements are key vehicles for resistance dissemination, particularly in contexts of environmental stress or high antimicrobial exposure^13^. While studies have linked gut mobilome profiles to clinical and ecological factors (e.g., hospital environments^14^, maternal transmission^15^), the influence of probiotic supplementation on the infant resistome and mobilome is largely unexplored, especially in full-term infants from LMICs. Similarly, the relationship between the gut resistome and fecal metabolome, which could enable its diagnostic use, is currently unknown.

Here, we leverage data from the ProRIDE randomized controlled trial (RCT), which enrolled 2,000 term-born, healthy newborns in rural Tanzania, to investigate the effects of a 4-week, home-administered multistrain probiotic on health outcomes and the gut microbiome^16^. In this study, we have examined a sub-cohort (*n* = 152, **Table S1**) comprising 80 infants who received probiotics and 72 who received a placebo, and whose fecal samples were characterized by i) deep shotgun metagenome sequencing with a spiked-in marine bacterium enabling absolute quantification, ii) culture-based detection of ESBL-E, and iii) targeted and untargeted metabolomics. We evaluated the impact of probiotics on the abundance and growth dynamics of AMR pathobionts, as well as the composition of the gut resistome, mobilome, and metabolome. This study provides novel insights into the mode of action of microbiome-targeted strategies for early-life AMR prevention and highlights the capacity of probiotics to shape the functional features of infants’ gut microbiome.

## MATERIAL & METHODS

### Ethics statement

Ethical approvals for the ProRIDE trial were obtained from the Regional Committee for Medical and Health Research Ethics in Western Norway (REK Vest 2019/1025), the Tanzanian National Institute of Medical Research (NIMR/HQ/R.8a/Vol.IX/3398), and the Tanzania Medicines and Medical Devices Authority (Authorization no.TZ21CT0002).

### Study design and participants

Design of the RCT, participant recruitment, and main results were described earlier^16,17^. Briefly, the trial was an investigator-initiated, single-site, double-blind, placebo-controlled RCT (1:1 ratio) conducted at Haydom Lutheran Hospital (HLH) and surrounding area in North-East Tanzania. A total of 2,000 healthy infants with a birth weight of ≥ 2.0 kg, born at HLH, other health facilities, or at home, were enrolled within the first three days of life and randomly assigned to receive either a multistrain probiotic (*n* = 1,000) or a placebo (*n* = 1,000). This study focuses on a subset of 152 participants (**Table S1**) with deep metagenomic sequencing of fecal samples collected at two time points: 6 weeks and 6 months of age (250 samples). Additionally, samples from this subset were analyzed using both untargeted (200 samples from 101 infants) and targeted (241 samples from 152 infants) metabolomics profiling.

The intervention group received Labinic® probiotic drops (Biofloratech Ltd, Surrey, UK). Each daily dose of five drops (0.2 ml) contained 2 billion colony-forming units (CFU) of *Lactobacillus acidophilus* NCFM, *Bifidobacterium bifidum* Bb-06, and *Bifidobacterium longum* subsp. *infantis* Bi-26, with equal amounts of all three strains. The placebo consisted of inactive ingredients. Both solutions were identical in taste, color, and packaging. Trained field workers conducted home visits at 1 week, 6 weeks, and 6 months of life. During these visits, a standardized electronic case report form was administered to collect data on recent illnesses, medication use, breastfeeding practices, growth, and the child’s overall condition. The field workers also collected a stool sample at both the 6-week and 6-month visits. The sample was transferred to eSwab® (Copan Diagnostics, CA, USA) and two collection tubes from DNA Genotek (Ottawa, Canada), which stabilize fecal DNA (OMNIgene•GUT | OM-200) and metabolites (OMNImet•GUT | ME-200). It was then transported in cold bags to HLH and stored at -80 [. The eSwab and the OMR-200 tubes were later transported on dry ice to Norway^16^.

The clinical data were collected from clinical records and caregiver interviews. For this sub-study specifically, the data included sex (male or female), hospitalization (defined as any inpatient admission, excluding the immediate post-delivery stay unless extended for medical reasons), place of birth (categorized as HLH, other health facility and home delivery), antibiotic exposure (assessed during follow-up visits at 6 weeks and 6 months post-enrollment using structured case report forms), and detection of ESBL-E carriage by culture.

### ESBL-E screening

Screening for ESBL-E was performed to evaluate the rates of ESBL-E gut colonization. Fecal samples collected with eSwab® were cultured using chromogenic agars selective for ESBL-E (ChromID ESBL, BioMérieux, Marcy l’Étoile, France). All samples were, in addition, plated on non-selective blood agar for growth control. Agar plates were incubated at 35°C in a normal atmosphere for 18-24 hours before inspection. In cases where no growth was observed on the blood agar, the sample was considered invalid. Samples with no growth on the chromogenic agars after 18-24 h of incubation were reported negative for ESBL-E. ESBL-E suspect colonies were identified at the species level using MALDI-TOF MS (Bruker Daltonik). An ESBL-E phenotype was confirmed using the double disk approximation test (Liofilchem, Roseto degli Abruzzi, Italy).

### DNA isolation, metagenomic sequencing, and bioinformatics preprocessing

AmbioGen, Stockholm, Sweden, performed total DNA extraction from fecal samples and metagenome sequencing using Nanoball DNA technology (**Supplementary file**). Before all downstream analyses, pair-end reads were checked for quality using FastQC v.0.11.9^18^. The adaptor sequences and low-quality reads were filtered using Trim Galore v.0.6.10^19^ with the default parameters. The human DNA contaminant sequences were discarded from all samples by filtering out the reads mapped against the human reference genome (GRCh38, downloaded from NCBI GenBank in 2022) using Bowtie2 v.2.4.4^20^ with -*-very-sensitive -- end-to-end* parameters. The identified paired reads that did not map against the human genome using SAMtools v.1.12^21^ with *-f 12 -F 256* were used in subsequent analyses.

### Metagenome, resistome, and mobilome profiling

Taxonomic profiling was obtained using MetaPhlAn4 v.4.0.6^22^ against the CHOCOphLAN database. The resistome annotation of metagenomic reads was performed by mapping them against the *nucleotide_fasta_protein_homolog_model* from the Comprehensive Antibiotic Resistance Database (CARD^23^ v.3.2.9) database using Bowtie2 v.2.4.4^20^ with parameter *– very-sensitive-local*. For the annotation of ARGs, a coverage threshold of 80% was used. Using SAMtools v.1.12^21^, the mapped reads were separated from the unmapped reads, sorted, and indexed, and the number of reads mapped for each ARG was calculated. The counts were then normalized for each sample to the total gene lengths by calculating reads per kilobase reference per million reads (RPKM). The clinical relevance of individual ARGs was based on the AMR Gene Family previously reported as clinically relevant by Diebold et.al.^24^ and the National Database of Antibiotic Resistant Organisms (NDARO). We excluded ARGs encoding efflux pumps and metabolic functions due to the difficulties of distinguishing their physiological roles from antibiotic resistance. The relative abundance of MGEs was quantified by employing ShortBRED v.0.9.5^25^. Specifically, MGEs sourced from the MobileOG v.1.6^26^ served as the proteins of interest for identifying marker families using ’*shortbred_identify.py’* with *’--clustid 0.95’* option. MGEs read counts were normalized in RPKM using ’*shortbred_quantify.py’*. Finally, we employed a computed peak-to-trough ratio (CoPTR) v.1.1.4^27^ to estimate the impact of probiotic supplementation on the replication rates of the two most important ESBL-E bacteria, *E. coli* and *K. pneumoniae*.

### Mass spectrometry (MS)-based metabolomics analyses

For both targeted and untargeted analyses, aliquots of fecal samples collected in the OMNImet•GUT were vortexed for 1 minute and centrifuged at 6,500 × *g* for 10 min at 4 °C. The supernatants were spin-filtered using a 10 kDa cutoff spin filter at 20,800 × *g* for 10 min at 0 °C. An aliquot of non-processed fecal sample (≈ 100 μL) was dried to determine the wet-to-dry weight ratio. This ratio was used to estimate theoretical dry weight of the samples analyzed. The metabolite levels were further adjusted for dilution and extraction volume. Final peak areas were normalized by dividing the final signal area by the estimated dry weight, yielding peak areas per gram of dried feces.

#### UPLC-MS/MS analysis for quantification of short-chain fatty acids and organic acids

For the targeted profiling of short-chain fatty acids (SCFAs) and organic acids (OAs), fecal samples were prepared according to a previously validated method^28^ with minor modifications. In brief, cleaned supernatants were derivatized in a 2:1:1 volume ratio with 200 mM 3-nitrophenylhydrazine (3NPH) and a 120 mM N-(3-dimethylaminopropyl)-N′-ethylcarbodiimide (EDC) solution (6% pyridine) prepared in a 50:50 (v/v) acetonitrile (ACN):MilliQ water mixture. The derivatization reaction was allowed to proceed for 30 minutes at 40 °C. The reaction mixture was then diluted first at 1:2 ratio with 2,6-di-tert-butyl-4-methylphenol (BHT) and subsequently in 1:2 ratio with 2% formic acid, both prepared in ACN:MilliQ (10:90, v/v), to quench the reaction. Lastly, the quenched mixture was diluted at 1:6.25 ratio with ACN:MilliQ (10:90, v/v), yielding a total dilution after derivatization of 1:25. Before MS analysis, the extract was mixed with an equal volume of isotopically labelled internal standards (ISTD_T).

#### Preparation of external and internal standards and quality controls

Analytical standards were purchased from Sigma-Aldrich (Darmstadt, Germany) and consisted of 15 SCFAs: acetate, propionate, butyrate, isobutyrate, valerate, isovalerate, pivalate, 2-methylbutyrate, methyl isobutyrate, hexanoate, 3,3-dimethylbutyrate, 2-ethylbutyrate, 2-methylvalerate, 3-methylvalerate, and 4-methylvalerate. Additionally, we used standards for eight OAs: lactate, succinate, pyruvate, oxaloacetate, α-ketoglutarate, fumarate, malate, and citrate. Analytical grade 3NPH, EDC, pyridine, formic acid, and BHT were also purchased from Sigma-Aldrich. For ISTD_T preparation, ^13^C-3NPH was purchased from Cayman Chemicals.

Preparation of external standard (ESTD) curves and ISTD_T for all targeted metabolites was performed as previously described^28^. In brief, 20 mM ESTD mixes of 1) acetate, propionate, and butyrate, 2) the remaining SCFA, and 3) all OAs, were freshly prepared in 50% ACN:MilliQ (v/v). The standard mixes were diluted to 5 mM for acetate, propionate, and butyrate, and one mM for the rest. This mixture was further diluted as an 8-point calibration curve with a two-fold dilution between the two highest concentrations, followed by a three-fold dilution between the other concentrations (3.43-5000 μM for acetate, propionate, and butyrate, and 0.69-1,000 μM for the rest). Because of much higher levels of lactate in the infant feces than any other metabolite, an additional standard mixture for lactate was prepared at a concentration of 30 mM and diluted to a 6-point curve, ranging from 937.5 to 30,000 μM, with a 2-fold dilution between each point. The two sets of standards were then derivatized and processed using the same concentrations and conditions as described for the fecal samples above. A fecal sample from an adult donor was processed in the same way as the standards/samples to serve as an internal quality control (QC). It was injected every 10 samples throughout the analytical run. ISTD_T mixes were prepared by diluting acetate to 4 mM, propionate and butyrate to 2 mM, and the rest of the metabolites to 1 mM. Fifty μL of each ISTD_T were mixed with 1 mg of 13C-3NPH, 25 μL EDC-6% pyridine, and 25 μL 50% ACN:MilliQ (v/v). The mixture was derivatized using the same conditions as above and, after adding BHT and formic acid, diluted to 100 ml in 10% CAN. Aliquots of the ISTD_T mixture containing all targeted metabolites were stored at -20 ^°^C for three months.

#### Targeted UPLC-MS/MS instrumentation

Targeted metabolomic analysis was conducted using Waters XEVO TQ-XS Triple Quadrupole mass spectrometer coupled to Waters Acquity UPLC. Chromatographic separation was achieved by gradient elution on Acquity Premier HSS T3 (1.8 μm, 2.1x100 mm). Mobile phase A consisted of MilliQ water with 0.1% formic acid (v/v), and mobile phase B of ACN with 0.1% formic acid (v/v). The gradient was as follows: 15% B for 2 min, 15-40% B in 10 min, 40-100% B in 1 min, 100% B for 0.1 min, and held at 15% B for 2 min. The flow rate was 0.5 mL/min, the column temperature 50 °C, and the autosampler was maintained at 6 °C. The mass spectrometer was operated in negative electrospray ionization mode using multiple reaction monitoring. Desolvation temperature was 550 °C, desolvation gas flow 1000 L/h, cone gas flow 150 L/h, nebulizer gas flow 7 bar, capillary voltage 0.6 kV, and ion energy 1 and 2 were set to 1 and 2, respectively. Peak integration and data processing were performed using the TargetLynx application in MassLynx (version 4.2 SCN 1007).

Analyte responses were calculated as a ratio of the endogenous peak area to corresponding ISTD_T peak area. Quantification was based on a weighted (1/x²) linear regression of the ESTD curve, using at least six calibration points that covered the biological concentration range, with an R² value close to 0.99. Two metabolite pairs, isovalerate/ methylisobutyrate and pivalate/ 2-methylbutyrate, did not separate and were thus quantified together.

#### Untargeted LC-MS/MS profiling

Supernatants from fecal samples were prepared as for the targeted analysis. Five μL of each sample supernatant was pooled to generate a QC sample. Afterwards, all supernatants including the QC were diluted 1:10 with ACN: MilliQ water (90:10, v/v) containing internal standard mix (ISTD_U) of Acetyl-l-carnitine-D_3_, Hexadecanoyl(palmitoyl)-l-carnitine-D_3_, l-Leucine-5,5,5-D_3_, l-Tryptophan-(indole-D_5_), l-Methionine-(methyl-D_3_), Stearic acid-18,18,18-D_3_, Chenodeoxycholic-2,2,3,4,4,6,6,7,8-D_9_ acid, 18:0-D_35_ Lyso PC, Dopamine-1,1,2,2-D_4_ hydrochloride (Merck Life Science, Darmstadt, Germany). Before sample analysis, the dilution factor, solvent type, sample extraction method, and sample stability in the instrument autosampler, freezer, and at room temperature were evaluated based on efficient sample handling, peak shape and intensity, as well as reproducibility, and found to be well within the dynamic range of the instrument. The final samples for analysis were randomized (RAND function in Excel, Microsoft Office) and transferred to 96-well injection plates (Waters, SKU: 186002481) with the QC sample every ten injections and kept in the autosampler (6 °C) throughout the acquisition.

#### LC-MS/MS instrumentation and data acquisition

Untargeted metabolomic profiling was based on a published hydrophilic interaction liquid chromatography (HILIC) method ^29^. In summary, the instrument was a Thermo Vanquish UHPLC system coupled with a Thermo Scientific Orbitrap ID-X Tribrid Mass Spectrometer. The data acquisition was performed using Thermo AcquireX Deep Scan module in data-dependent acquisition (DDA). Chromatographic separation was achieved by injecting 3 μL of each sample through a BEH Amide (100x2.1 mm, 1.7 Å) column at a flow rate of 0.45 mL/min. The column temperature was maintained at 50 °C, and the autosampler at 6 °C. The separation was carried out using gradient elution with mobile phase A consisting of H_2_O with 10 mM NH_4_Ac at pH 9, and mobile phase B of ACN: H_2_O (9:1, v/v) with 10 mM NH_4_Ac at pH 9. Mobile phase B was maintained at 100% for 6 min to equilibrate the system and then kept at 100% for an additional 2.5 min after injection. Then 100-60% from 2.5-9 min and kept at 60% for 0.2 min.

The ion source was operated in heated negative electrospray ionization (2500 V). Gas settings were 50 arbitrary units (arb) for sheath gas, 10 arb for auxiliary gas, and one arb for sweep gas, with an ion transfer tube temperature of 325 °C and vaporizer temperature of 350 °C. Full MS scans were acquired using the Orbitrap at a resolution of 120,000 over a scan range of 70-800 m/z. The automatic gain control (AGC) target was set to custom with a normalized AGC of 25 and a maximum injection time of 50 milliseconds (ms). For MS/MS, up to 8 dependent scans were triggered per MS1 event using higher-energy collision dissociation (HCD, with stepped collision energies of 20%, 35%, and 50%) and collision-induced dissociation (CID, with a fixed energy of 30%). MS2 scans were also acquired in the Orbitrap at a resolution of 30,000 with a quadrupole isolation window of 1.2 m/z and a maximum injection time of 150 ms. Dynamic exclusion was enabled (excluding after one acquisition, for 5 seconds) with isotope exclusion to prevent repeated fragmentation of isotopologues.

#### Untargeted LC-MS/MS quality control assessment

Prior to sample injection, a system blank consisting of clean sample solvent (ACN: MilliQ, 90:10, v/v) was injected to ensure that the instrument and mobile phase were free from contamination. Next, several injections of a system control sample consisting of 0.1 mg/L quercitin, amitriptylin, histidine, arginine, labetalol, doxepin, proline, and tryptophan (Merck Life Science, Darmstadt, Germany) were injected to monitor and ensure instrument performance, *e.g*., retention time, mass accuracy, peak shape, and peak intensity. When all the above quality checks were acceptable, a processed and diluted clean solvent from OMNImet•GUT | ME-200 tubes was injected twice as an extraction blank. Then, four injections with the pooled QC were injected to generate deep scan ID samples before unknown samples were run in full scan mode. The pooled QC was injected every 10 samples to monitor drift along the run.

Additionally, the ISTD_U spiked into every sample was also used to monitor the drift and quality of each injection. After the injection sequence, the ISTD_U m/z values were uploaded into the software Skyline (v 24.1.0.414, MacCross Lab, University of Washington, USA), where retention time drift and signal intensity were monitored for each ISTD_U in each sample. Next, extracted ion chromatograms were manually checked and compared in the software Freestyle (Thermo Fisher Scientific, San Jose, USA). Specific masses corresponding to tryptophan, proline, and isoleucine were manually verified in each of the QCs to confirm peak shape, retention time, and intensity. The blanks were also checked to verify the absence of the specific masses. Finally, to evaluate stability, QCs were visualized in a PCA plot to detect potential global drift in metabolomic profiles over time.

#### Raw MS/MS data processing

The raw data files were centroided and processed using MZmine 4.7.8^30^ in two processing batches. The processing workflows were as follows: in the first batch, mass detection in MS1 was performed for all samples using a noise level of 1.0e4 based on precursor intensity. Then, mass detection in MS2 was performed without noise level filtering for four pooled MS2 samples, which were representative of the MS2 data. Subsequent steps included chromatogram builder, smoothing (Savitzky-Golay), local minimum feature resolver, 13C isotope filter, and isotopic peaks finder with default settings. Processing batch two was performed by running the join aligner first on the merged ID samples, then aligning to all the full scan samples using a retention time tolerance of 0.2 min. The extraction blank was used in the module feature list blank subtraction with a fold change threshold of 300% to retain relevant features. The following modules were applied: group MS2 scans with features, feature list rows filter, correlation grouping, and ion identity networking. The processed data were extracted for downstream analysis in GNPS^31^ and SIRIUS^32^. To determine the need for drift correction, the quantification table was uploaded to MetaboAnalyst 6.0, and the QCs were monitored using principal component analysis (PCA).

#### SIRIUS-based annotation

For feature annotation, the processed data from MZmine were uploaded into SIRIUS 6.2.2 ^32^ using default settings. Spectral matching, molecular formula prediction, CSI: FingerID^33^, and structure database search with PubChem as a fallback was enabled. Annotation was manually curated and evaluated based on the compound confidence score. A score above 0.6 was considered good if the substructure matched with databases and had a consistent top structure hit. These features were assigned a level 2 annotation. Features with lower confidence scores, which showed reliable substructure matches, were assigned level 3 annotation and were annotated at the ClassyFire^34^ most specific class level.

#### Feature-based molecular networking and statistics

For feature-based molecular networking, the processed data from MZmine were uploaded and run in GNPS2^31^. The processed network was then uploaded to the software Cytoscape 3.10.3^35^ for network visualization. Edges were labeled with their delta mz and further colored by cosine (blue) or MS1 annotation (red). Furthermore, the node sizes were mapped to the sum of precursor intensity. The normalized quantification table was uploaded to MetaboAnalyst, filtered by standard deviation (40%), normalized by median, log-transformed, and Pareto scaled. To assess significant features, a fold change cutoff of 2, with a p-value threshold of 0.05, and a false discovery rate (FDR) were used, along with unequal group variance, to generate a list of significantly different features between the two groups. This list was further uploaded to Cytoscape. The nodes were colored either for significantly higher (red) or lower (blue) abundance to visualize global trends in the network.

### Statistical analyses

#### Linear mixed models and differential abundance analyses

The α-diversity analysis was performed using *vegan* v.2.5.7^36^ R package. We evaluate the statistical influence of variables on the diversity and relative abundance of the infant microbiota, resistome, and mobilome using linear mixed-effects modelling (LMM) with the *lmerTest* v.3.1.3^37^ R package with ‘infant_id’ as a random effect. To standardize the resistome and mobilome relative abundance, we applied an inverse-normal transformation (*qnorm* function in R). Additionally, to detect collinearity among the variables, we used a variance inflation factor (≤ 5) using the *car* v.3.1.3^38^ R package. We used the *emmeans* v.1.10.7^39^ R package to compute estimated marginal means following LMM to evaluate the effect of covariates at each time point. Linear regression models with the Benjamini-Hochberg procedure for *p*-value adjustment were employed to identify differentially abundant SCFAs, considering ‘trial_arm’, ‘sex’, ‘ESBL-E’, ‘born_hospital’, and ‘antibiotics’ as covariates. The *ANCOM-BC* v.2.2.2^40^ R package with the Benjamini-Hochberg procedure for *p*-value adjustment was used to identify differentially abundant features at the species level. In this model, we considered ‘trial_arm’, ‘sex’, ‘ESBL-E’, ‘born_hospital’, and ‘antibiotics’ as fixed effects. Differential abundance was considered statistically significant at *p*[<[0.05 and *q*[<[0.25, given the high dimensionality of the data. We employed MaAsLin 3 v.0.99.4^41^ R package using *normalization = ‘NONE’*, *median_comparison_abundance = FALSE*, and *median_comparison_prevalence = FALSE* to identify species-level differential abundance based on absolute abundances normalized using a spike-in bacteria count^42^, as follows:

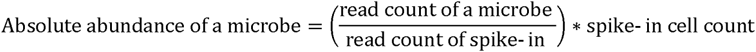

The read count of a microbe was based on MetaPhlAn4, and the read count of the spike[in microbe was 1.4x10^5^ cells of *Vibrio campbellii*. In the MaAsLin model, we also considered ‘trial_arm’, ‘sex’, ‘ESBL-E’, ‘born_hospital’, and ‘antibiotics’ as fixed effects.

#### ***β***-diversity, ordination, and clustering analyses

β-diversity analysis was executed using the *“vegdist”* function from the *vegan* v.2.5.7^36^ R package, applying Bray-Curtis or Jaccard dissimilarity indices, followed by principal coordinate analysis (PCoA) for ordination. To ascertain the statistical significance of composition differences, we conducted a permutational multivariate analysis of variance (PERMANOVA) using the *“adonis”* function from the *vegan* package, with 999 permutations. The Redundancy Discriminant Analysis (RDA) was performed using *vegan* v.2.5.7^36^ R package. The microbiota clusters were generated using an unsupervised non-linear k-means partitioning algorithm based on *Bifidobacterium* genus relative abundance using the *kmean*s function in base R. Ordination was performed using non-metric multidimensional scaling (NMDS) based on Bray-Curtis dissimilarity using *vegan* v.2.5.7^36^ R package. The significance and effect size of each covariate were evaluated using the *’envfit’* function based on 999 permutations from the *vegan* v.2.5.7^36^ R package.

#### Other statistical analyses and data visualization

All statistical analyses were conducted using R software v.4.4.1. Graphical illustrations were mostly created with the *ggplot2* v.3.3.6 R package^43^. The heatmaps were visualized using *ComplexHeatmap* v.2.20.0 package^44^. Random forest was performed using 10-fold cross-validation, 500 trees, and 1000 permutations using *randomForest* v.4.7.1.2^45^ and *caret* v.6.0.94^46^ R packages to identify important predictors of ESBL-E carriage. We employed a generalized linear model (GLM) with a binomial family using the *glm* function in R to evaluate the impact of various factors on the carriage of ESBL-E. Additionally, we fitted negative binomial models using the *MASS* v.7.3.65^47^ R package. We employed Spearman’s correlation analysis using *Hmisc* v.5.1.3^48^ R package to investigate the relationships between microbiota, resistome, and mobilome diversities, as well as the correlation between the relative abundance of *B. longum* and the richness of potential pathogens. We performed partial least squares path models using *plspm* v.0.5.1^49^ R package to identify pathways between microbial composition, ARGs, and MGEs, allowing the quantification of direct and indirect effects between these components. Pathway coefficient β > 0.3 was considered a strong interaction. We performed DIABLO (Data Integration Analysis for Biomarker Discovery using Latent Variable Approaches for Omics Studies) from the *mixOmics* v.6.28.0^50^ R package to explore associations between microbiota, resistome, and metabolome, aiming to identify key features that contribute to the relationship between these datasets. Subsequently, a biweight midcorrelation analysis was conducted using WGCNA v.1.73^51^ R package to assess the strength and robustness of associations between the selected features identified by DIABLO. Statistical assumptions were evaluated using the *AssumpSure* v1.0.0^52^ R package to ensure transparency in test selection. Relevant diagnostic figures are provided in the supplementary “**Assumptions**” file.

#### Statistical analyses of metabolomic data

Dry-weight normalized untargeted metabolomic data were uploaded to MetaboAnalyst 6.0^53^ and filtered using a low-variance filter based on a standard deviation threshold of 40%. The resulting data were median log_10_-transformed and Pareto scaled. Significant features between groups were determined by a Mann-Whitney U test (p < 0.05) for unequal group variance and with FDR correction and visualized as volcano plots with a fold change cutoff of 2.0 and a *p*-value threshold of 0.05. PCA was carried out using the Euclidean distance matrix, and PERMANOVA for statistical testing. For targeted metabolomics, the relative abundance of metabolites was visualized using the R package *ggplot2*, and correlation heatmaps were generated using Spearman correlation with a *p*-value threshold of 0.05.

## RESULTS

### Stability of Probiotic Species Colonization in the Infant Gut

For this sub-study of the ProRIDE trial, we selected 152 infants whose samples underwent thorough metagenomics and metabolomics analyses (**Table S2**). Probiotic supplementation during the first 4 weeks of life significantly reshaped gut bacterial composition, with notable effects evident at 6 weeks of age, but not at 6 months (**Figure 1a, Table S3**). At 6 weeks, the probiotic group had significantly higher relative abundances of *B. longum* and *B. bifidum* compared to the placebo group (ANCOM-BC2, Benjamini-Hochberg adjusted *p* < 0.01; **Figure 1b; Table S4**). In addition to these two *Bifidobacterium* species found in the probiotic product, we detected a significant increase in *Morganella morganii*, a species from *Enterobacterales* order in the probiotic group. Although *Lactobacillus acidophilus* was among the supplemented probiotic species, it was not detected in the metagenomic data.

**Figure 1.**
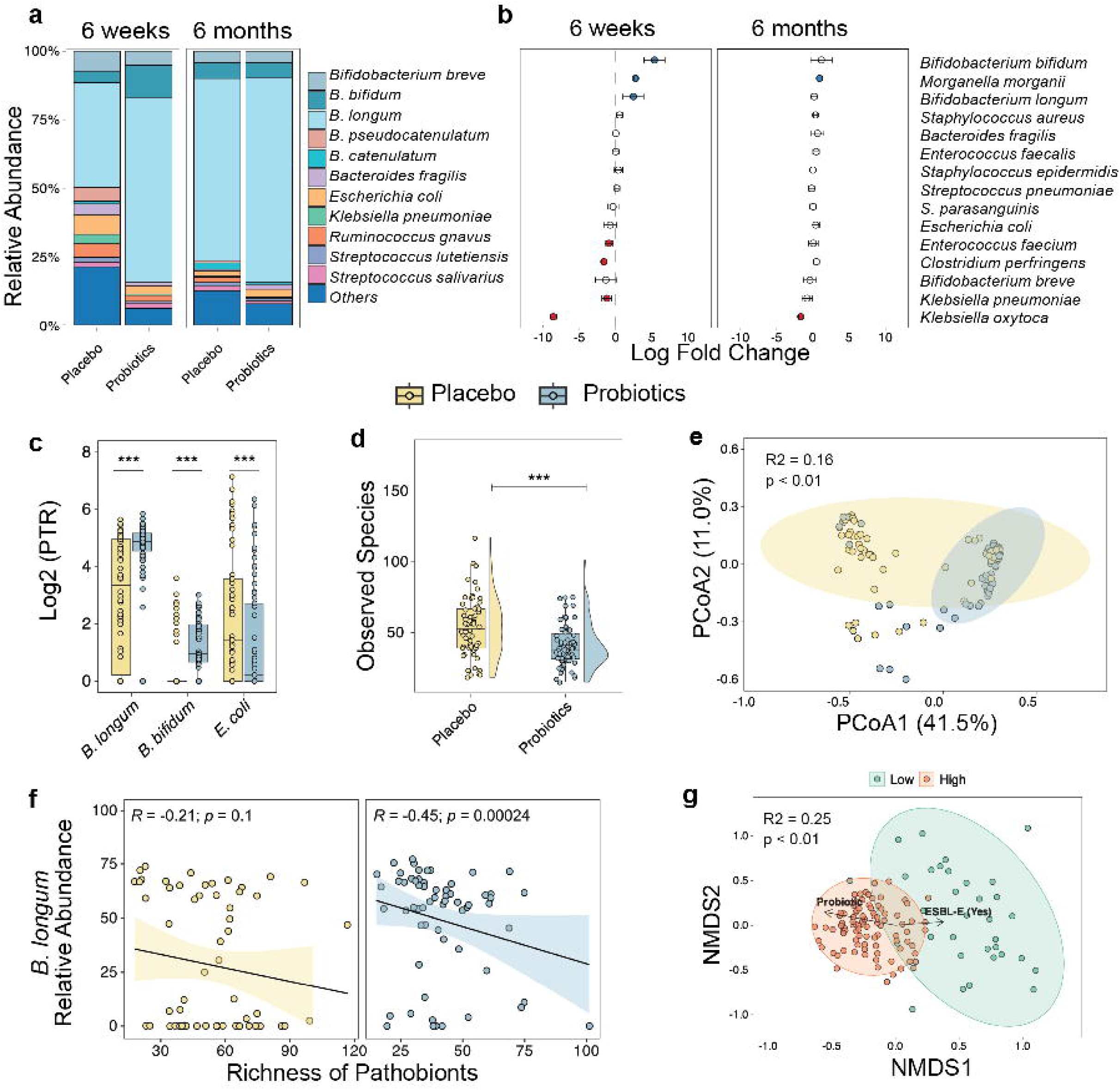
Impact of Oral Probiotics on Bacterial Composition of Infant Gut. (a) Relative abundance of bacterial species in placebo and probiotic-supplemented groups, inferred by MetaPhlAn4. (b) Differential abundance of species with pathogenic potential, identified using ANCOM-BC2 (Benjamini-Hochberg adjusted *p*-values). (c) Replication rate analysis was performed using the Compute Peak-to-Trough Ratio (CoPTR). *p*-values were computed using the Mann-Whitney U test. (d) Comparison of observed species richness between groups, with each point representing a sample. *p*-values were computed using a negative binomial model. (e) Principal Coordinate Analysis (PCoA) illustrating microbiota beta diversity differences. (f) Spearman correlation analysis between *B. longum* relative abundance and the richness of pathobionts. (g) Non-metric multidimensional scaling (NMDS) and PERMANOVA illustrate dissimilarities between clusters, with a biplot showing correlations between variables and clusters. Statistical significance: *p* < 0.05. Symbols: ∗∗∗ *p* < 0.001; ∗ *p* < 0.05.

Conversely, the relative abundance of *K. pneumoniae* was significantly lower in the probiotic group at 6 weeks (ANCOM-BC2 model; Benjamini-Hochberg adj. *p* = 0.014). Probiotic supplementation also significantly reduced the abundance of other bacteria with pathogenic potential: *Enterococcus faecium, Klebsiella oxytoca*, and *Clostridium perfringens* (ANCOM-BC2; Benjamini-Hochberg adjusted *p* < 0.05, **Figure 1b**). Most of the observed taxonomic shifts were transient and not present at 6 months, except for a persisting trend of lower abundance of *Klebsiella oxytoca* and increased abundance of *Morganella morganii* in the probiotic group (**Tables S5).**

To validate these findings, we utilized absolute abundance data derived from a unique quantification methodology employed prior to the sequencing process. Quantitative measurements of bacterial abundance were enabled by spiking the fecal samples with DNA of a known amount from the marine bacterium *Vibrio campbellii* (**Tables S6)**. To determine differences between the probiotic and placebo groups, we applied MaAsLin 3^41^, which permits the assessment of absolute abundances, in contrast to ANCOM-BC2^40^, which can only be used on relative data. Consistently, the cell counts of *B. longum* and *B. bifidum* were significantly higher in the probiotic group than in the placebo group at 6 weeks, confirming probiotic species colonization upon supplementation (model accounting for sex, trial arm, place of birth, and antibiotic exposure Benjamini-Hochberg adjusted *p* < 0.01; **Table S7**). In contrast to the relative abundance data, absolute abundances were not significantly different for any other species. At 6 months, we did not detect any significant differences in bacterial species abundances based on the absolute data (**Table S8)**.

We next inferred microbial growth dynamics from the metagenomic data by performing replication rate analysis using the CoPTR tool, which calculates the ratio of sequencing coverage at the replication origin to that at the replication terminus in a bacterial species^27^ (**Table S9**). We found that probiotic supplementation significantly increased the growth rates of *B. longum* and *B. bifidum* (Mann-Whitney U test, *p* < 0.0001) while decreasing the growth of *E. coli* at the 6-week time point (Mann-Whitney U test, *p* < 0.029; **Figure 1c**).

Lastly, we assessed α- and β-diversity of the bacterial communities to determine the broader impact of probiotics on gut microbiota. The α-diversity analysis revealed that the probiotic group had significantly lower bacterial richness compared to the placebo group at 6 weeks (negative binomial model; *p* < 0.01; **Figure 1d** & **Figure S1a).** Ordination analysis based on Bray-Curtis distances showed distinct microbial communities between the probiotic and placebo groups at 6 weeks (PERMANOVA: Bray-Curtis distance; R² = 16.11%; 999 permutations; *p* < 0.01; **Figure 1e**) but not at six months (**Figure S1b)**. We also observed that the probiotic group had a significantly less diverse microbiota composition than the placebo group at the 6-week timepoint (**Figure S1c**; Mann-Whitney U test, *p* < 0.01). Although we observed no significant differences in β-diversity at 6 months, significant differences in the Shannon index persisted for up to 6 months, indicating lasting effects of early-life probiotic use (**Figure S1d**; Mann-Whitney U test, *p* < 0.01).

### Probiotic-Mediated Reduction of Pathobionts

Building on the finding of reduced abundance of pathobiont species by probiotics, we investigated the association between the relative abundance of *B. longum* and the richness of pathobionts. We selected 18 bacterial species with established pathogenic potential (**Table S10**)^54–56^. Spearman correlation analysis revealed a moderate negative correlation between the relative abundance of *B. longum* and the richness of pathobionts, which was significant in the probiotic-supplemented group at 6 weeks (R=-0.45; *p* < 0.01; **Figure 1f**).

As a follow-up, we applied unsupervised clustering of metagenomic samples to identify co-occurring bacteria. This analysis, which used k-means clustering based on the abundance of the *Bifidobacterium* genus, identified two clusters: “Low” characterized by lower abundance of *Bifidobacterium* spp. and higher abundance of pathobionts, and “High” defined by higher *Bifidobacterium* spp. and lower pathobiont abundance (**Figure S2a**). The probiotic group was significantly more likely to belong to the “High” cluster compared to the placebo group at 6 weeks (*p* < 0.01, Fisher’s exact test, **Figure S2b**).

To identify factors correlated with the bacterial composition of the two clusters, we performed the Envfit analysis on the NMDS ordination (**Figure S2c)**. The tested variables included trial arm, sex, hospitalization, place of birth, antibiotic use, and detection of ESBL-E carriage by culture (**Table S2**). We were unable to test the impact of feeding type because the vast majority of the included infants were breastfed (148/152; 97%). At 6 weeks, probiotic supplementation emerged as the strongest explanatory factor of the microbiota composition (R² = 0.194; *p* < 0.01), followed by ESBL-E carriage (R² = 0.086; *p* < 0.01). By 6 months, the rate of ESBL-E carriage was the only significant factor associated with the clusters’ composition (R² = 0.084; *p* < 0.01; **Figure S2d**).

Consistent with these findings, NMDS based on Bray-Curtis dissimilarity and PERMANOVA showed clear separation of samples into “Low” and “High” clusters (R² = 0.25; *p* < 0.01, **Figure 1g**). Probiotic supplementation was strongly associated with the “High” cluster, while ESBL-E carriage was linked to the “Low” cluster, and their obtuse angle in the ordination plot indicated a negative correlation between these factors. These results were validated by a generalized linear model (GLM), which showed a significant negative association between probiotic supplementation and ESBL-E carriage at 6 weeks (*p* < 0.001; **Figure S3a**). A random forest model further identified probiotic supplementation as the most important predictor of ESBL-E carriage at 6 weeks (**Figure S3b**).

### Impact of Probiotics on the Gut Resistome and Mobilome

Next, we assessed the impact of probiotic supplementation on the resistome. We analyzed metagenomic data against the CARD database, focusing on resistome profiles independently of microbiota composition. We initially identified 1,214 ARGs, which were filtered to 919 more clinically relevant ones that confer resistance to 15 antibiotic classes (**Table S11**). At 6 weeks, dominant ARGs in the placebo group were linked to multidrug resistance (*i.e*., conferring resistance to two or more antibiotic drugs), tetracyclines, aminoglycosides, beta-lactams, and diaminopyrimidines (trimethoprim) (**Figure 2a**). In the probiotic group, the distribution of the ARG classes was similar to the placebo group at both time points, but their relative abundance was significantly lower at 6 weeks of age (Mann-Whitney U test; *p* < 0.01; **Figure 2b**).

**Figure 2.**
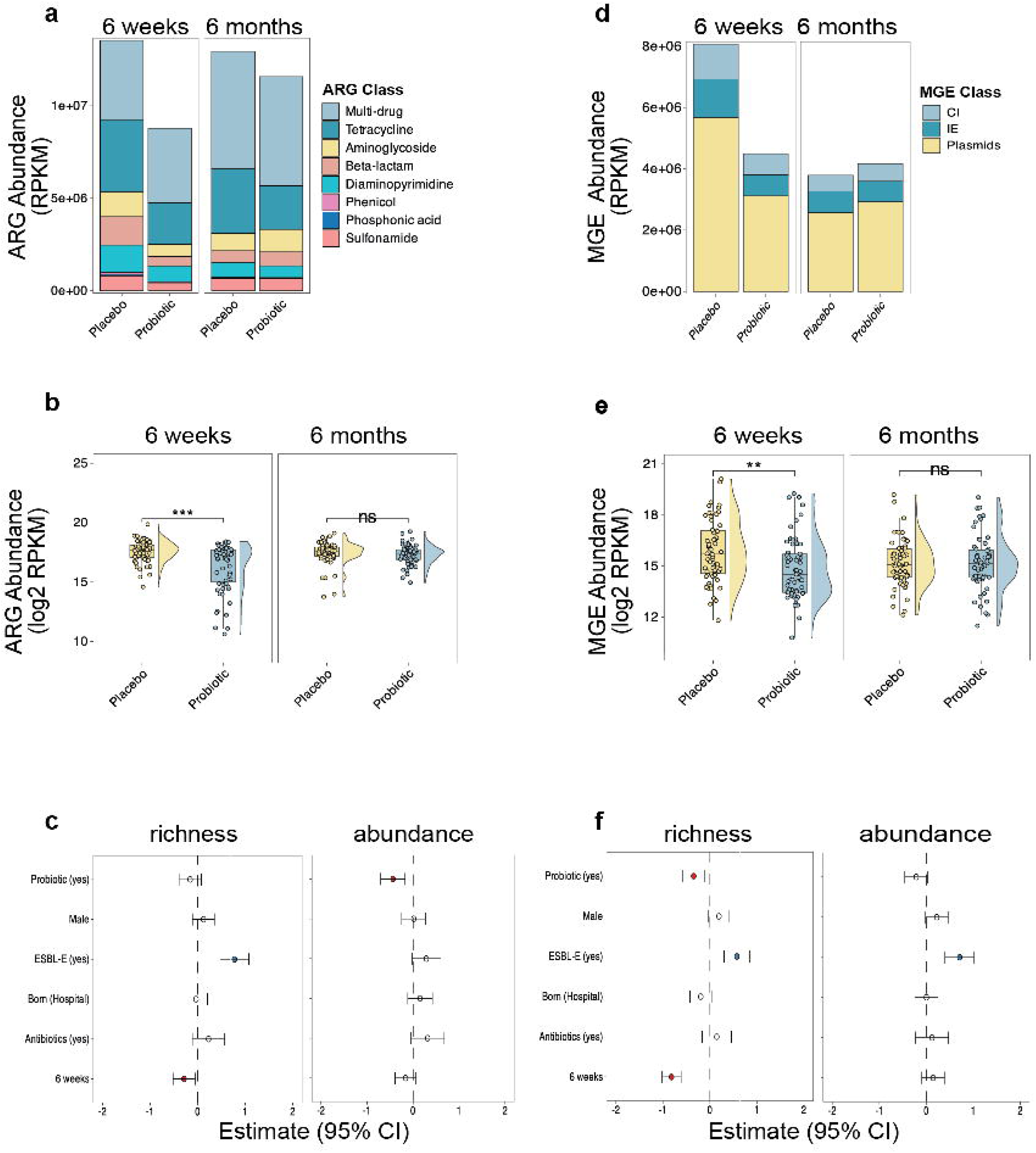
Impact of Probiotics on the Gut Resistome and Mobilome. (a) Relative abundance of antibiotic resistance genes (ARGs) in reads per kilobase per million mapped reads (RPKM), stratified by ARG classes. Multidrug-resistant class denotes ARGs conferring resistance to two or more antibiotic drugs. (b) Comparison of ARG relative abundance between groups, with each point representing a sample. (c) The impact of selected variables on the resistome richness estimated by a negative binomial model and relative abundance as determined by linear mixed-effect modelling (LMM). (d) Relative abundance of mobile genetic elements (MGEs) in RPKM, stratified by MGE classes. (e) Comparison of MGE relative abundance between groups, with each point representing a sample. (f) The impact of selected variables on the mobilome richness estimated by a negative binomial model and relative abundance as determined by LMM. Statistical significance: *p* < 0.05. *p*-values were computed using the Mann-Whitney U test. Symbols: ∗∗∗ *p* < 0.001; ∗∗ *p* < 0.01.

LMM analysis revealed that ESBL-E carriage was significantly associated with an increased ARG richness, while probiotic supplementation was associated with decreased ARG abundance (LMM; *p* < 0.001, **Figure 2c**). Estimated marginal means (EMMs) analysis of probiotic impact on resistome richness and relative abundance at 6 weeks of age, further validated this effect (*p* = 0.0316 and *p* < 0.001, respectively, **Figure S4a**). However, we did not detect any change in resistome richness or abundance between the groups at the six-month time point, suggesting that the effect of probiotic supplementation during the first month of life did not persist until six months. On the other hand, when we extended the resistome analysis to the “Low” and “High” clusters, the “Low” cluster had a higher resistome relative abundance than the “High” cluster (Mann-Whitney U test; *p* = 0.0261; **Figure S4b**). Lastly, ordination analysis using Jaccard distances revealed a distinct resistome composition between the probiotic and placebo groups at 6 weeks (PERMANOVA: R² = 7.23%; 999 permutations; p < 0.01), but this difference was no longer present at 6 months (**Figure S4c)**.

The mobilome facilitates the horizontal transfer of ARGs between bacterial cells. Therefore, we analyzed the mobilome by querying the metagenomic data against the MobileOG database^26^ (**Table S12**). Plasmids were the most abundant MGEs (**Figure 2d**), and the mobilome relative abundance was significantly lower in the probiotic group compared to the placebo group at six weeks of age (Mann-Whitney U test; *p* < 0.01; **Figure 2e**).

LMM analysis showed that the carriage of ESBL-E was significantly associated with both mobilome richness and load (LMM; *p* < 0.001; **Figure 2f**). Complementary analysis of EMMs revealed a significant reduction in mobilome richness and load at six weeks of age (*p* < 0.01 and *p* < 0.02, respectively; **Figures S5**). Finally, consistent with resistome findings, β-diversity analysis using Jaccard distances showed a distinct mobilome composition between probiotic and placebo groups at six weeks (PERMANOVA: R² = 2.5%; 999 permutations; *p* < 0.01). None of these associations was present at six months of age.

### Interactions between Bacterial Microbiota, ARGs, and MGEs

Considering the similarities between microbiota, resistome, and mobilome profiles in the context of probiotic supplementation, we sought to determine the extent to which bacterial community structure influences ARG and MGE profiles. Spearman correlation analysis identified moderate to strong correlations between α-diversities of microbiota, resistome, and mobilome (**Figure S6**). To describe interaction pathways between bacterial composition, ARGs, and MGEs, we employed partial least squares path modelling. We assumed that the microbiota affects both resistome and mobilome, and estimated the correlation between resistome and mobilome. In the first model (**Figure 3a**), which estimated the influence of mobilome on resistome, bacterial microbiota had a strong direct effect on MGEs (path coefficient β = 0.79, *p* < 0.01) and a weak direct effect on ARG composition (β = 0.10, *p* < 0.01). At the same time, microbiota exhibited a strong indirect impact on ARGs via MGEs (β = 0.56, *p* < 0.01), and MGEs had a strong direct effect on ARG composition (β = 0.71, p < 0.01). To explore whether ARGs also impact MGE composition, we modified the model (**Figure 3b**) by allowing resistome data to predict interactions with mobilome. This adjustment revealed a strong direct effect of ARGs on MGEs (β = 0.46, *p* < 0.01). In this model, microbiota was a strong predictor of both MGEs (β = 0.49, *p* < 0.01) and ARGs (β = 0.66, *p* < 0.01) and had a moderate indirect effect on MGEs through ARGs (β = 0.30, *p* < 0.01). Altogether, these models can be interpreted as follows: (1) microbiota is likely to influence ARG composition through MGEs, and (2) MGE composition contributes to ARG profile, reinforcing their interconnectedness.

**Figure 3.**
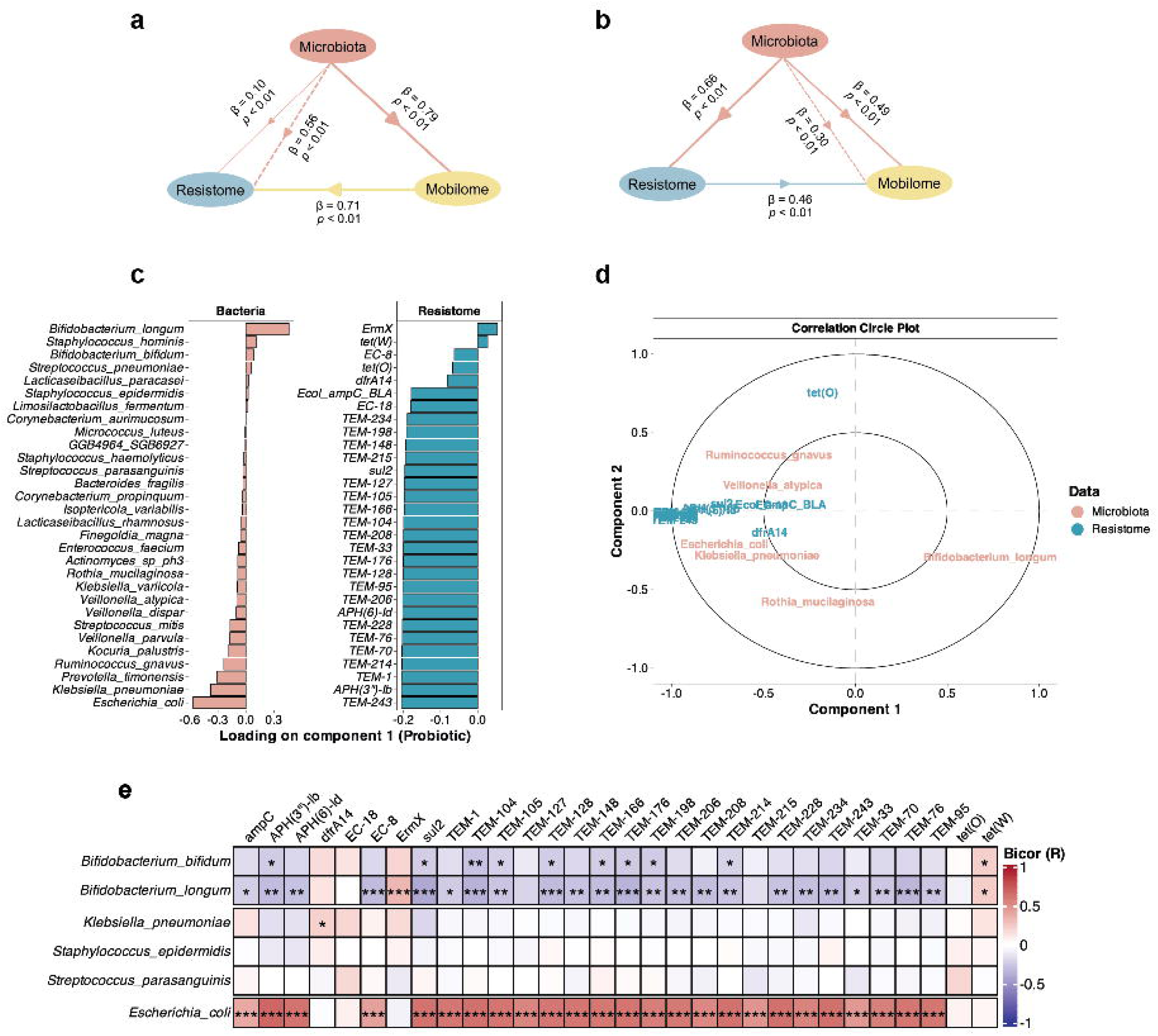
Interactions Between Microbiota, Resistome, and Mobilome. Partial least squares path modelling illustrates interactions between microbiota, resistome, and mobilome, with (a) model 1 set to use mobilome data to predict interactions with resistome; (b) model 2 using resistome data to predict interactions with mobilome. (c) Top 30 most variable predictors identified by DIABLO. (d) Correlation circle plot illustrating relationships between bacterial species and ARGs. (e) Correlation between selected pathobionts and probiotic species, determined using biweight midcorrelation. All analyses are based on the microbiome data of 6-week samples. Statistical significance: *p* < 0.05. Symbols: ∗ *p* < 0.05; ∗∗ *p* < 0.01; ∗∗∗ *p* < 0.001

We then predicted bacterial hosts of ARGs by data integration analysis for biomarker discovery using latent components (DIABLO). Among the top 30 discriminative predictors, we observed that the relative abundance of several pathobionts and ARGs was associated with the placebo group (**Figure 3c**). In contrast, commensal bacteria, such as *B. longum*, were associated with the probiotic group. Additionally, the correlation circle plot revealed strong negative correlations between *B. longum* and key pathobionts, *E. coli,* and *K. pneumoniae* (**Figure 3d**), as well as between *B. longum* and the most variable ARGs. In contrast, *E. coli* and *K. pneumoniae* exhibited strong positive correlations with the resistome, consistent with our previous findings, which describe them as major ARG reservoirs^57,58^. To validate these predictions, we performed a biweight midcorrelation analysis (Bicor) and filtered the results to retain 18 selected bacterial species with pathogenic potential^54–56^ and probiotic species.

Features with low variance or missing values that resulted in undefined correlations were excluded from the analysis. The results aligned with the DIABLO output: *E. coli* showed a strong positive correlation with the top 30 most variable ARGs, whereas *B. longum* had a strong negative correlation (**Figure 3e**).

### Impact of Probiotic Supplementation on Fecal Metabolome

We continue our investigations by investigating microbiota’s functional output, *i.e*., fecal metabolite profiles, by applying untargeted metabolomics to stool samples from 101 infants, comprising 200 samples (**Table S11**). Similarly to the metagenomic data, we observed no significant differences in untargeted metabolomic data between the probiotic and placebo groups in the 6-month-old infants. However, PCA of metabolite profiles derived from 6-week samples showed distinct clusters for probiotic and placebo groups (Euclidean distance, PERMANOVA, R 0.07, *p* value 0.001, **Figure S7a**). We detected 45 significantly increased and 33 decreased metabolic features when comparing the groups at 6 weeks (Mann-Whitney U test, p < 0.05, FDR correction, **Figure S7b)**. Most of these were putatively annotated to belong to carbohydrates and amino acids, but without clear clustering on feature-based molecular networking. Moreover, when we compared the 6-week samples based on their *Bifidobacterium* spp. abundance, the “High” and “Low” groups formed well-separated groups (Euclidean distance, PERMANOVA, R 0.25, *p* value 0.001, **Figure 4a**). Samples from infants that belonged to the “High” cluster showed 190 significantly increased and 354 decreased metabolite features (Mann-Whitney U test, p<0.05, FDR, **Figure 4b**), supporting a broad metabolic impact of early-life *Bifidobacterium* colonization.

**Figure 4.**
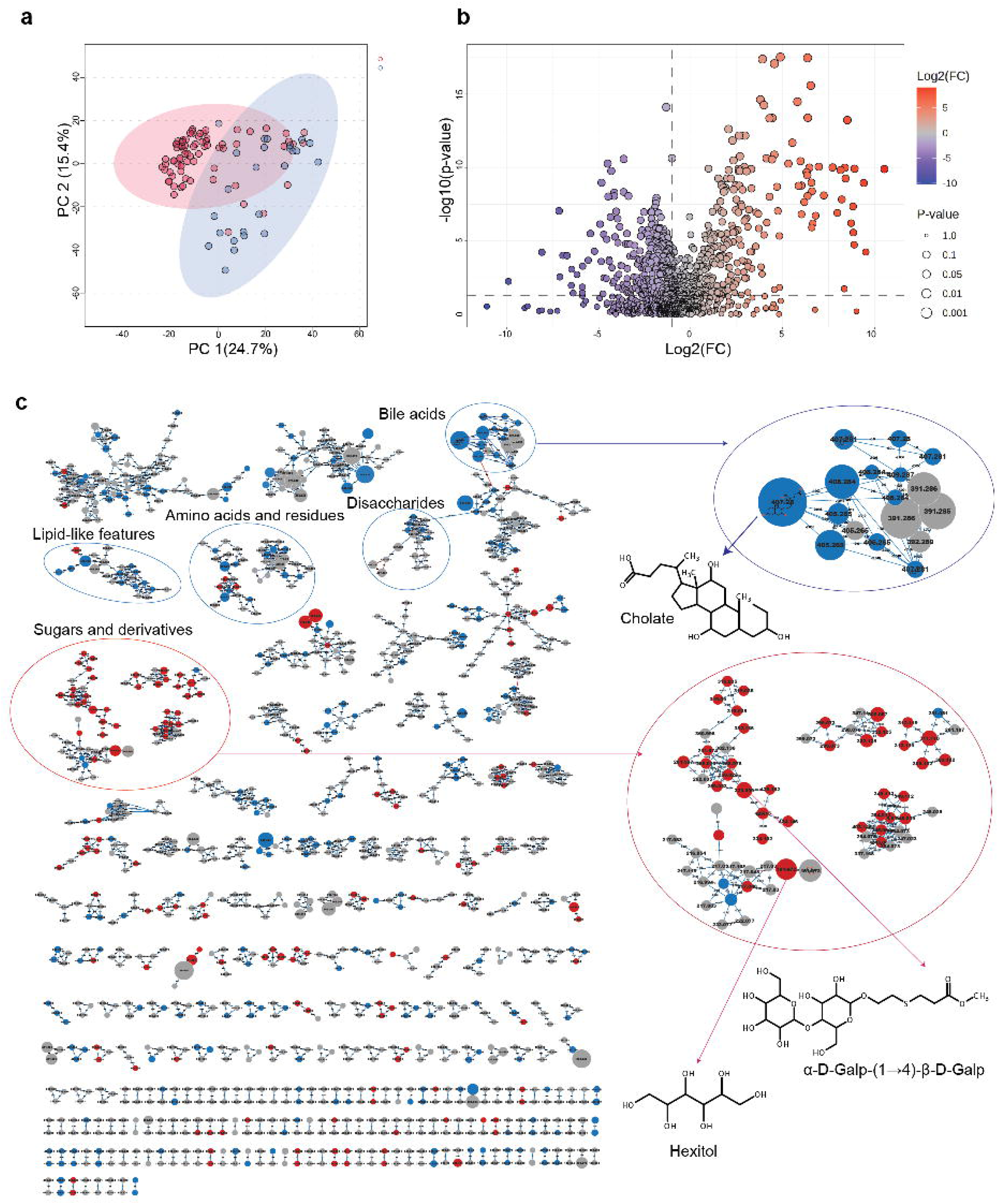
Untargeted metabolomics of infant stool samples from 6 weeks. A) Principal component analysis of “High” (red) and “Low” (blue) clusters based on *Bifidobacterium* spp. abundance (Euclidean distance, PERMANOVA, R = 0.251, p = 0.001). B) statistical comparison of the “High” and “Low” clusters’ metabolic profiles visualised as a Volcano plot (Two-sided unpaired Student’s t-test, *p*<0.05, FDR, 2-fold change threshold, and unequal group variance). In the “High” cluster, there were 354 and 190 features significantly decreased and increased, respectively, as compared to the “Low” cluster. C) Feature-based molecular networking showing clustering based on cosine (blue edge) and MS1 (red edge) similarities between features. MS1 refers to using the initial mass spectra data, *i.e*., information on the mass-to-charge ratio and intensity of ions, to identify and characterize metabolites. Node size reflects the sum of the precursor area for each feature labelled by its ion mass.

Closer examination revealed that metabolites with significantly increased abundance in the “High” group were putatively annotated to belong to simple sugars and derivatives, where hexitol and a disaccharide containing two galactopyranoses were annotated with high confidence (confidence score 0.518 and 0.955, respectively). On the other hand, metabolite features with significantly decreased abundance were putatively annotated as belonging to lipid-like compounds, amino acid residues, di- and oligosaccharides, and bile acids (**Figure 4c**). Notably, the bile acid cholate was annotated with high confidence based on spectral matching (confidence score 0.976). Cholate and structurally related metabolites also had one of the highest abundances in the metabolite profile. These results link probiotic supplementation to alterations in both microbial and host metabolites.

### Microbial Metabolites Predictive of ESBL-E Carriage

Considering the impact of probiotics on the fecal metabolome, we inquired whether specific microbial metabolites play a role in the identified pathobiont suppression and decreased ARG load. Using targeted metabolomics, we quantified 23 metabolites (15 SCFAs and 8 Oas, **Table S13**), which are key microbial metabolic intermediates and end products. A comparison of these 23 metabolites in 241 stool samples from 152 infants (**Table S2**) revealed significantly higher levels of citrate, pyruvate, and lactate in the probiotic group, whereas propionate levels were significantly higher in the placebo group at 6 weeks (linear regression model with the Benjamini-Hochberg correction, **Figure 5a**). In addition, several SCFAs (isobutyrate, isovalerate/methyl isobutyrate, and propionate) were positively associated with ESBL-E carriage, while pyruvate, acetate, and lactate showed negative associations.

**Figure 5.**
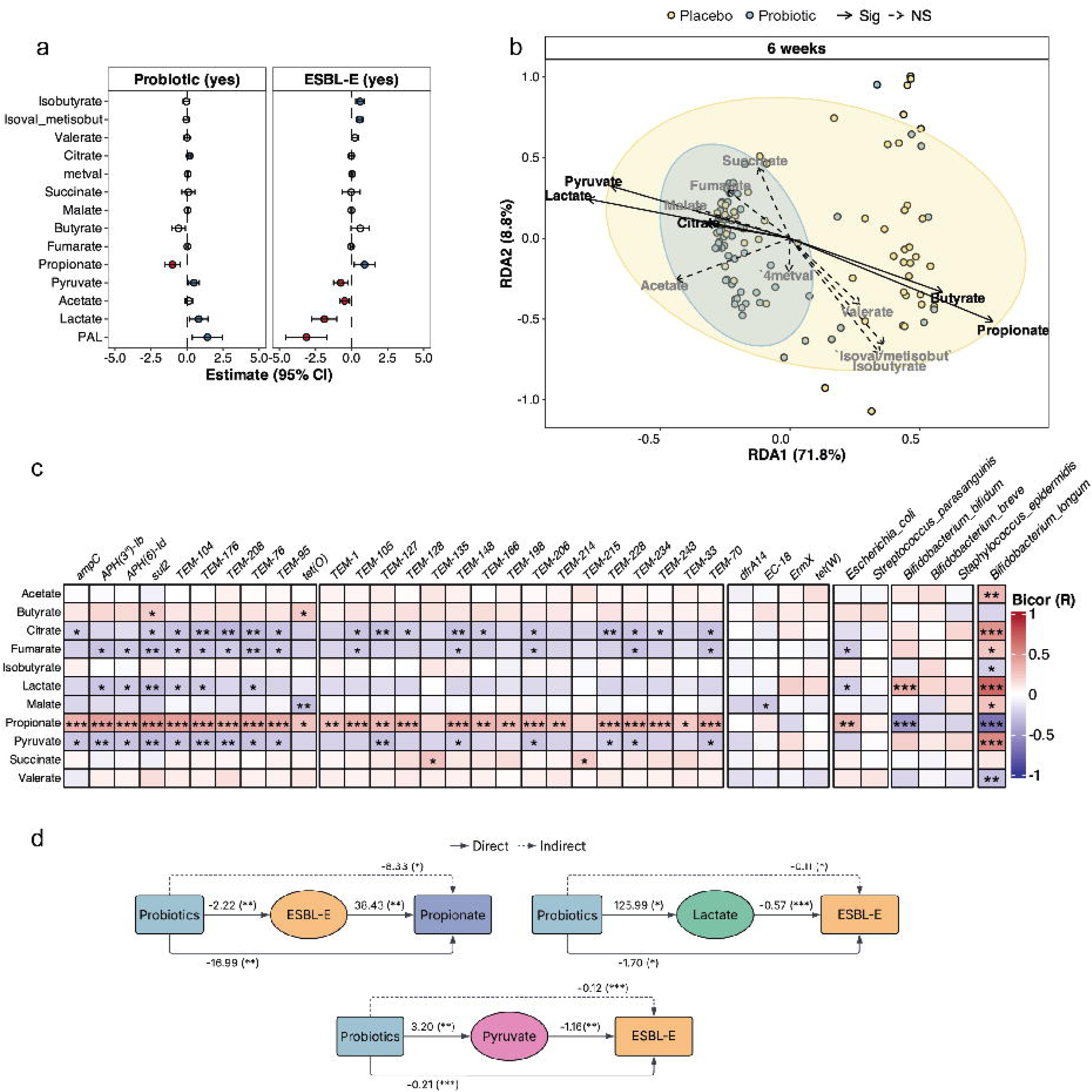
Targeted metabolomics of infant stool samples from 6 weeks. (a) Differential abundance of the metabolites identified using linear model (Benjamini-Hochberg adjusted p values). PAL – combined output of pyruvate, acetate, and lactate. (b) Redundancy discriminant analysis biplot depicting the relationship between the bacterial communities and short-chain fatty acids (SCFAs). (c) Biweight midcorrelation analysis showing correlations between metabolites and the top 30 most variable ARGs. (d) Three different models based on the mediation analysis testing the effect of probiotics on Extended-spectrum beta-lactamase-producing Enterobacterales (ESBL-E) colonization. Statistical significance: *p* < 0.05. Symbols: ∗ *p* < 0.05; ∗∗ *p* < 0.01; ∗∗∗ *p* < 0.001

In infancy, primary fermenters such as *Bifidobacterium* spp. mainly produce acetate and lactate through the Bifid shunt, *i.e*., fructose-6-phosphate pathway^59^, whereas secondary fermenters produce propionate and butyrate most often through cross-feeding from metabolites such as lactate, acetate, and succinate^60^. To closely examine metabolites involved in the Bifid shunt through which bifidobacteria metabolize hexose sugars, we combined the absolute abundances of pyruvate, acetate, and lactate. We identified that this metabolite combination further increases their positive association with the probiotic group. Extending the analysis to clusters based on *Bifidobacterium* spp. abundance, the levels of lactate, pyruvate, butyrate, and propionate profiles were also significantly different between the “High” and “Low” clusters (**Figure S8a**).

To investigate the relationship between SCFAs and OAs and overall microbial community structure, we performed redundancy discriminant analysis. This analysis showed that citrate, pyruvate, and lactate were significantly associated with microbiota composition in the probiotic group. In contrast, propionate and butyrate were significantly associated with microbial profiles in the placebo group (**Figure 5b**). These associations were supported by a random forest model, where propionate, lactate, citrate, and pyruvate emerged as the top predictors of microbiota composition (**Figure S8b**). Finally, by using DIABLO, we validated that lactate and pyruvate are the top predictors of the probiotic group, while propionate levels predicted the placebo group (**Figure S8c**). These associations were supported by biweight midcorrelation analysis. Specifically, acetate, citrate, lactate, and pyruvate were significantly associated with *B. longum*, while propionate was significantly correlated with *E. coli*.

Additionally, propionate showed a strong positive correlation with many of the top 30 variable ARGs, particularly *bla*_TEM_, whereas citrate, fumarate, lactate, and pyruvate were significantly and negatively associated with these ARGs (**Figure 5c**).

Propionate, pyruvate, and lactate were associated with both probiotic supplementation and ESBL-E carriage, suggesting that these metabolites lie on the mechanistic path linking probiotics and reduced ESBL-E carriage. To test this hypothesis, we performed a mediation analysis, testing the hypothesis that probiotics affect ESBL-E. We found that probiotics potentially reduce propionate levels indirectly by decreasing the ESBL-E carriage (**Figure 5d**). Additionally, the analysis indicated that lactate and pyruvate directly mediate the effect of probiotics on ESBL-E, *i.e*., the probiotic strains reduce ESBL-E carriage through increased levels of these metabolites.

### Reinforcement of *Bifidobacterium* Metabolic Signatures by Probiotic Supplementation

To further describe how the probiotic strains affect the metabolite dynamics, we examined indicators of metabolic fluxes, that is, metabolite ratios and correlations of the individual SCFAs and OAs. At 6 weeks, infants with “High” *Bifidobacterium* spp. abundance, in both the probiotic and placebo groups, showed a consistent dominance of lactate and acetate, with lower levels of other SCFA and OAs such as succinate (**Figure S9a**). In contrast, infants with “Low” *Bifidobacterium* abundance displayed a more diverse metabolic profile, with higher proportions of succinate, propionate, and butyrate compared to the “High” groups. The latter results suggest an increased activity of secondary fermenters that rely on cross-feeding.

In the “High” *Bifidobacterium* groups, lactate and acetate showed significant positive correlation (probiotic group - Spearman coefficient R=0.66; p < 0.001; placebo group - R=0.18; p < 0.001, **Figure S9b**), consistent with co-production of these metabolites via the Bifid shunt pathway. On the other hand, the 6-week placebo “Low” *Bifidobacterium* group showed a significant negative correlation between succinate and propionate (R=-0.47; p < 0.01, **Figure S9c**), indicating active conversion via the succinate pathway, a major route for microbial propionate production^61^ . This metabolite flux was not significant in the other groups.

Additional differences emerged when comparing samples from the placebo and probiotic groups. The placebo “High” *Bifidobacterium* group showed significant negative correlations between several SCFAs (propionate, butyrate, isobutyrate, valerate, and isovalerate/methyl isobutyrate) and OAs (citrate, fumarate, pyruvate, malate, succinate, and lactate), suggesting SCFAs production via these OAs intermediates. This pattern was less pronounced in the probiotic “High” *Bifidobacterium* group (**Figure S9b**), despite relatively similar metabolite abundances.

## DISCUSSION

In this study, we examined the impact of a 4-week probiotic supplementation in healthy full-term infants in Tanzania on the gut carriage of ESBL-E, growth rates of *E. coli* and *K. pneumoniae*, pathobiont richness, and the functional features of the gut microbiome. To develop a deep understanding of the probiotic mode of action, we explored the gut resistome and mobilome composition, their correlations with the structure of the gut microbiota, and fecal metabolome. Simply put, this study offers a comprehensive understanding of how probiotic bacteria can mitigate antibiotic resistance in high-risk populations.

We found that probiotics reduced the overall microbiota diversity, the relative abundance of pathobionts, and the replication rate of one of the key bacterial hosts of ARGs, *Escherichia coli*. Previous studies also suggested that probiotics can inhibit bacterial species with pathogenic potential^62–64^, including *E. coli* and *K. pneumoniae,* in the gut by competing for nutrients and adhesion sites. Additionally, probiotics can secrete bacteriocins, hydrogen peroxide, and other antimicrobial compounds that inhibit the growth of pathogenic bacteria^63^. Beyond direct antagonism, probiotics may enhance host immune responses, promoting the production of anti-inflammatory cytokines, which may further suppress pathogen colonization^64^.

Random Forest analysis showed that birth in the hospital and antibiotic exposure were key factors contributing to ESBL-E carriage, consistent with previous studies^65^. However, GLM analysis did not find these associations to be statistically significant. This may be due to the time gap between birth or antibiotic exposure and sample collection. The first samples were collected six weeks after birth, during which the influence of hospital exposure may have diminished, in line with previous findings^66^. Similarly, the effect of antibiotics is transient, with ARG carriage often declining after treatment cessation^67,68^. Beyond hospitalization and antibiotic use, ESBL-E carriage is shaped by broader environmental factors, including water quality, sanitation, and hygiene conditions, and environmental contamination^69^, highlighting the complex drivers of ESBL-E spread.

Our study found that probiotic supplementation significantly reduced resistome load, aligning with previous studies^11,70^. However, this effect was transient, as almost no differences were observed at the six-month time point. These results are consistent with prior findings showing that probiotic supplementation induces an initial but short-lived reduction in ARG burden^71^. Similarly, a study on hospitalized infants reported that probiotics lowered resistome diversity and prevented the persistence of ARGs in the gut microbiota up to five months of age^11^. The transient nature of this effect highlights the possibility that continuous or repeated probiotic supplementation may be required to sustain its beneficial impact on resistome load.

Probiotic supplementation reduced both the richness and load of MGEs, potentially limiting the horizontal transfer of ARGs between bacterial species. To our knowledge, this study is the first one that has explored the impact of probiotics on the infant gut mobilome, offering novel insights into how probiotics influence mobilome composition and its role in antibiotic resistance transmission. Additionally, we observed moderate to strong correlations between microbiota, resistome, and mobilome, consistent with previous studies^15,57^. Notably, *E. coli* emerged as a major ARG reservoir, emphasizing its role in antibiotic resistance transmission, consistent with previous studies^15,57,72–74^. The reduction in ESBL-E carriage and other pathobionts likely contributed to the diminished resistome and mobilome in the infant gut, as the resistome and mobilome are inherently linked to bacterial hosts. By suppressing key bacterial reservoirs, probiotics may indirectly limit ARG and MGE abundance and dissemination.

The results of partial least squares path modelling indicated that the relationship between microbiota, resistome, and mobilome is convoluted. The gut microbiota composition appeared to indirectly influence the resistome through the mobilome and, conversely, affect the mobilome through resistome alterations. The mobilome likely plays a central role in resistome expansion, as indicated also by a previous study^12^. Although not directly comparable, several *in vitro* studies have shown that ARG carriage can affect bacterial fitness and mobilome dynamics. For example, a single copy of *mcr-1*, a colistin-resistance gene, reduces the fitness of *E. coli*^75^. The persistence of *mcr-1* depends on plasmid copy number regulation, where *TrfA1*, a plasmid replication initiator protein, enhances plasmid replication, thereby promoting mobilome expansion and increasing horizontal gene transmission rates^76^.

By profiling the fecal metabolome, our study brought a nuanced understanding of the probiotic strain’s mode of action. The finding of a higher abundance of simple sugars concurrently with a lower abundance of di- and oligosaccharides confirms the role of *Bifidobacterium* spp. as primary fermenters degrading complex carbohydrates to simple sugars and fermentation end products. Our findings also align with an earlier report on probiotic supplementation in preterm infants, supporting the notion that *Bifidobacterium* spp. provide a favorable metabolic gut milieu with potentially improved fatty acid absorption^77^.

To get a better understanding of the probiotic-induced changes in fermentative metabolism, we determine absolute levels of SCFAs and OAs in the fecal samples. As predicted by a theoretical modeling study^59^, we identified high levels of pyruvate, acetate, and lactate in the 6-week samples from infants in the “High” *Bifidobacterium* groups. This metabolic pattern corroborates a gut ecosystem dominated by *Bifidobacterium* spp., which are primary fermenters that metabolize human milk oligosaccharides by the Bifid shunt pathway into acetate and lactate via pyruvate. Sugar derivatives such as simple organic acids may lower the intestinal pH, thus suppressing colonization of bacterial taxa that do not tolerate low pH, including drug-resistant pathobionts. We tested this hypothesis by mediation analysis, which suggested that lactate and pyruvate directly mediate the effect of probiotics on ESBL-E.

Although the placebo and probiotic groups both featured taxonomically similar “Low” and “High” *Bifidobacterium* groups, differences emerged when comparing their metabolite profiles. The placebo “High” *Bifidobacterium* group showed significant negative correlations between SCFAs and OAs, suggesting SCFA production via the organic acid intermediates.

This pattern was less pronounced in the probiotic “High” *Bifidobacterium* group, implying that probiotic supplementation results in reduced cross-feeding dynamics, likely resulting from the probiotic *Bifidobacterium* spp. dominance. Moreover, the more diverse metabolic profile of the “Low” *Bifidobacterium* group suggested increased activity of secondary fermenters that rely on cross-feeding to produce SCFAs such as propionate and butyrate. This phenomenon was particularly pronounced in the 6-week placebo “Low” *Bifidobacterium* group, with a metabolic profile indicating active conversion of succinate to propionate.

The relationship between ESBL-E carriage and SCFA has so far been mainly studied *in vitro*^78^ or in animal models^79^. One of the few human studies examined asymptomatic gut carriage of ESBL-producing bacteria in Dutch adults^80^. This study did not detect any link between ESBL-E carriage and fecal metabolites; however, the authors used nuclear magnetic resonance, which is less sensitive than LC-MS. An earlier study made similar observations to ours, but in preterm infants, where probiotic administration led to higher fecal acetate and lower pH in the supplemented infants^81^. Another study, which investigated longitudinal SCFA profiles from 12 children during the first 2 years, showed that infant microbiomes dominated by *Enterobacterales* were associated with reduced acetate and increased succinate^82^. Lastly, we determined a strong association between propionate, resistome, and ESBL-E carriage. In this context, findings from an *in vitro* study showing that propionate-containing medium stimulates virulent properties of pathogenic *E. coli*^83^, suggest that propionate might be beneficial for the proliferation of drug-resistant pathobionts.

We acknowledge several limitations in this study, including the limited number of time points for sample collection and the resulting constrained resolution of microbial dynamics.

Additionally, detailed data on antibiotic use, including type, duration, indication, timing after birth, and maternal intrapartum exposure, were unavailable, limiting our ability to assess their full impact. Similarly, dietary factors, such as the introduction of solid foods, which can influence microbiome composition, were also not extensively documented. Finally, the absolute abundance data derived from MetaPhlAn should be interpreted with caution because the tool does not provide real read counts for each bacterium, but only for their marker genes.

Despite these limitations, this study provides novel insights into how probiotic supplementation affects the infant gut microbiota, resistome, mobilome, and metabolome. Notably, we document that probiotics reduce ESBL-E carriage, resistome, and mobilome load by dominating the gut microbiota and metabolically altering the infant gut. Our findings highlight the importance of going beyond taxonomical composition and abundance to develop an understanding of functional relationships between microbial metabolites and their producers. And while probiotics show promise in lowering ESBL-E carriage in early life, the observed transient effects suggest that sustained supplementation or complementary strategies may be needed for lasting benefits. Future research should focus on optimizing probiotic formulations, dosing, and examining long-term impacts on microbiota stability and ARG transmission. Unravelling these mechanisms will be key to designing targeted microbiome interventions to combat AMR in vulnerable populations.

## Supporting information

Suppmenetary file

## CONTRIBUTORS

VKP designed this sub-study of the ProRIDE trial, with contributions from CK and AB. NL, CK, IHL, and VKP secured funding. MJ, SJM, and BB performed data curation. AB crafted the bioinformatics pipeline, conducted the analyses, performed statistical and machine learning analyses, multi-omics integration, and data visualization. AB also drafted the initial manuscript. GHB carried out all metabolomics experiments and data analyses. VKP supervised data analyses and manuscript preparation and finalized the figures. All authors provided critical feedback on the manuscript and approved the final version for submission.

## DECLARATION OF INTERESTS

The authors declare that they have no competing interests.

## FUNDING

This trial received main funding from Western Norway Regional Health Authority (grant no. 912267), Trond Mohn Foundation (TMS2020TMT11), JPIAMR STRESST project (NFR333432), and Northern Norway Regional Health Authority (grant no. HNF-1512-20 and HNF1705-24). Additional funding was provided by Centre for New Antibacterial Strategies (CANS) through Tromsø Research Foundation to VKP (TFS18_CANS_AS-HVF).

## DATA SHARING STATEMENT

The shotgun metagenomic reads, with human DNA removed, have been deposited to SRA under BioProject ID PRJNA1305341. The identified marker families from the MobileOG database (v.1.6) are available here (https://doi.org/10.5281/zenodo.11491353). The untargeted metabolomics will also be publicly available in the MassIVE repository (https://massive.ucsd.edu, accession ID MSV000098821) after the peer review.

## ACKNOWLEDGEMENTS

We extend our heartfelt thanks to the field workers at Haydom Global Health Research Centre for their dedication and hard work. Special gratitude is extended to Ketil Størdal, Blandina Theophil Mmbaga, Charles Makasi, and Sven Gudmund Hinderaker for their oversight of the Data and Safety Monitoring Board; Thomas Eagan for providing randomization support; and Helene Heitmann Sandnes for managing the randomization plan. We also thank AmbioGen for DNA extraction and sequencing, as well as Sigma2 Norwegian Research Infrastructure Service for their computing resources. Above all, we are deeply grateful to the village leaders and families around Haydom Lutheran Hospital for their trust and participation; this study would not have been possible without them.

## Notes

### Competing Interest Statement

The authors have declared no competing interest.

