## Supplementary material for "Metabolic reprogramming of the infant gut by bifidobacteria-based probiotics drives exclusion of antibiotic-resistant pathobionts": Suppmenetary file

Bargheet et al.

**CONTENT**

**DNA extraction and sequencing**

**Figure S1**. **α- and β-diversity analyses**

**Figure S2**. **Unsupervised clustering and Envfit analyses.**

**Figure S3**. **Impact of probiotics on ESBL-E carriage**

**Figure S4.** **The influence of probiotics on the resistome**

**Figure S5.** **The influence of probiotics on the mobilome.**

**Figure S6. Correlation analysis.**

**Figure S7. Comparison of the fecal metabolome of placebo and probiotic groups at 6 weeks**

**Figure S8 Targeted metabolomics data analysis.**

**Figure S9.** **Dynamics of SCFAs and OAs based on *Bifidobacterium* spp. abundance**

**DNA extraction and sequencing**

Fecal samples were thawed and mixed with liquefaction reagent (DNA Genotek, Canada). To each sample, 10 µL of a *Vibrio campbellii* culture (10⁴ cells/µL) was added as an internal spike-in control. A total of 800 µL of the mixture was transferred to ZR BashingBead lysis tubes containing a combination of 0.1- and 0.5-mm beads (Zymo Research, USA) for mechanical lysis. Each extraction batch included a positive and a negative control: the positive control consisted of 75 µL of the ZymoBIOMICS Microbial Community Standard (Zymo Research, USA) in 725 µL liquefaction reagent, while the negative control consisted of 700 µL liquefaction reagent and 100 µL of *Vibrio campbellii* culture (10⁴ cells/µL).

Samples underwent mechanical lysis using the FastPrep-24 5G instrument (MP Biomedicals, USA) and were subsequently centrifuged at 10,000 × g for 1 minute. A volume of 200 µL of the resulting supernatant was transferred to a deep-well plate for DNA extraction. The extraction was performed using the Tecan Fluent liquid handling platform (Tecan, Switzerland) following the ZymoBIOMICS 96 MagBead DNA protocol (Zymo Research). DNA was eluted in 50 µL of EB buffer (Qiagen, Germany) and stored at –20 °C until further use.

Following DNA extraction, samples were quantified using the QuantIT High Sensitivity dsDNA Assay on a Tecan Spark plate reader (Tecan, Switzerland). Samples with DNA concentrations above 1.1 ng/µL were spiked with 1% *Alicyclobacillus acidophilus* DNA (relative to total input), while samples with concentrations below 1.1 ng/µL were supplemented with *Alicyclobacillus* DNA to reach a total of 50 ng per reaction. The negative sequencing control was also spiked with *Alicyclobacillus* DNA to monitor potential background signal.

Library preparation was carried out using 50 ng of input DNA and the MGI FS Library Prep Set (MGI Tech, China), following the manufacturer's protocol. Library quality was assessed using the Agilent TapeStation D1000 kit (Agilent Technologies, USA), and concentrations were confirmed using the same QuantIT assay. Equimolar amounts of the resulting libraries were pooled to a final concentration of 100 pM, circularized using the MGI Easy Circularization Kit (MGI Tech, China), and sequenced on the DNBSEQ-T7 platform (MGI Tech, China) using 150 bp paired-end reads, according to the manufacturer's instructions.

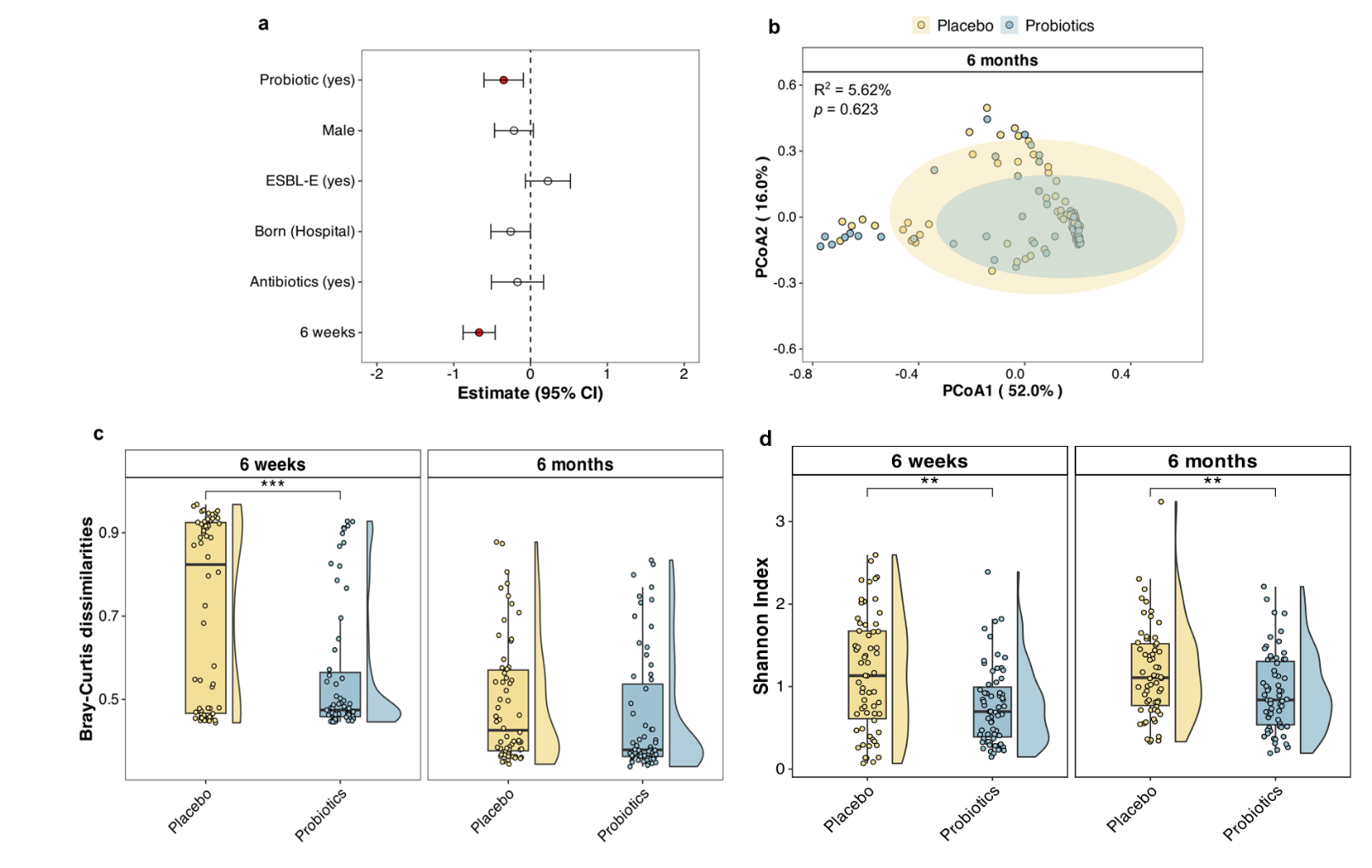
**Figure S1**. **α- and β-diversity analyses (**a) Impact of selected variables on infant gut microbiota richness, determined by a negative binomial model. (b) Principal Coordinate Analysis (PCoA) illustrating microbiota beta diversity differences at 6 months. Changes in beta and alpha diversities between the probiotic and placebo groups, visualised through comparisons of (c) Bray-Curtis dissimilarity, and (d) Shannon index (Mann-Whitney U test). The Wilcoxon effect size for 6 weeks = 0.276, magnitude small; for 6 months, effect size = 0.233, magnitude small. Statistically significant results are based on *p* < 0.05. *** *p* < 0.001, ** *p* < 0.01.

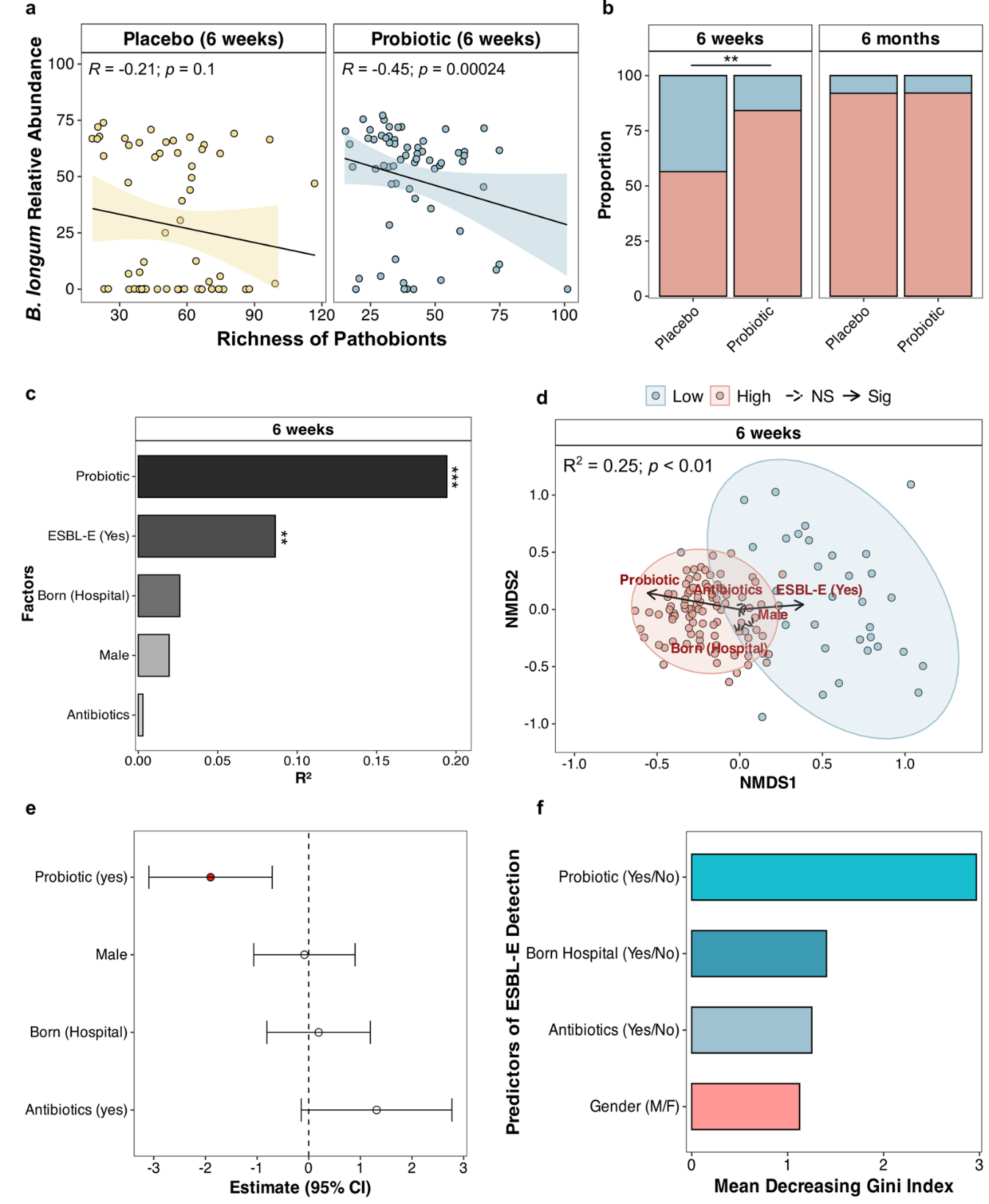
**Figure S2**. **Unsupervised clustering and Envfit analyses.** a) Microbiota cluster composition identified using k-means clustering, with Cluster 1 ("Low") and Cluster 2 ("High"). (b) Distribution of clusters across groups, with the *p*-value calculated using Fisher's exact test. Factors associated with microbiota community structure, identified using Envfit analysis, at 6 weeks (c) and 6 months (d) of age. Statistical significance: *p* < 0.05. Symbols: ∗∗∗  *p* < 0.001; ∗∗  *p* < 0.01; ∗ *p* < 0.05.

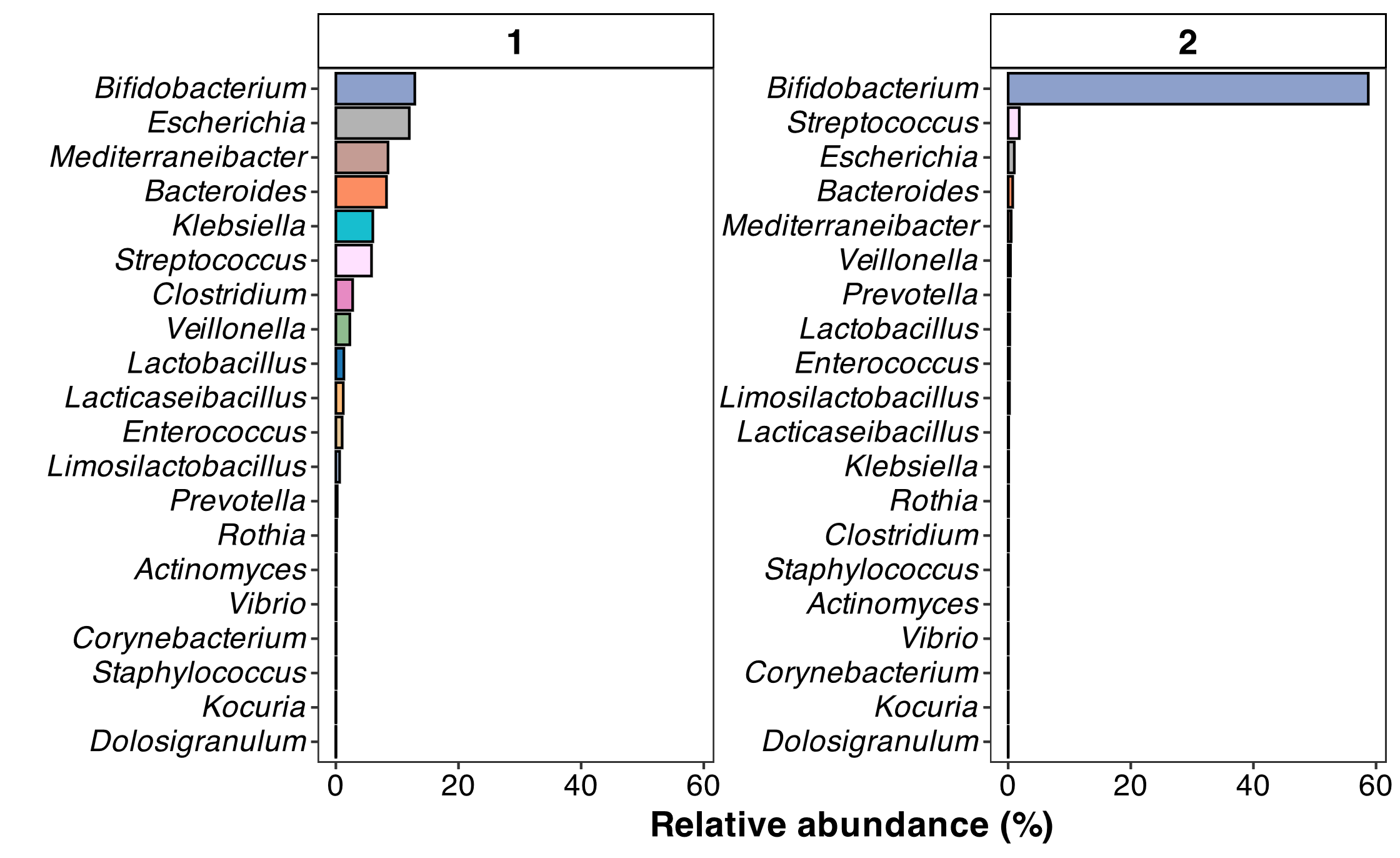

**a**

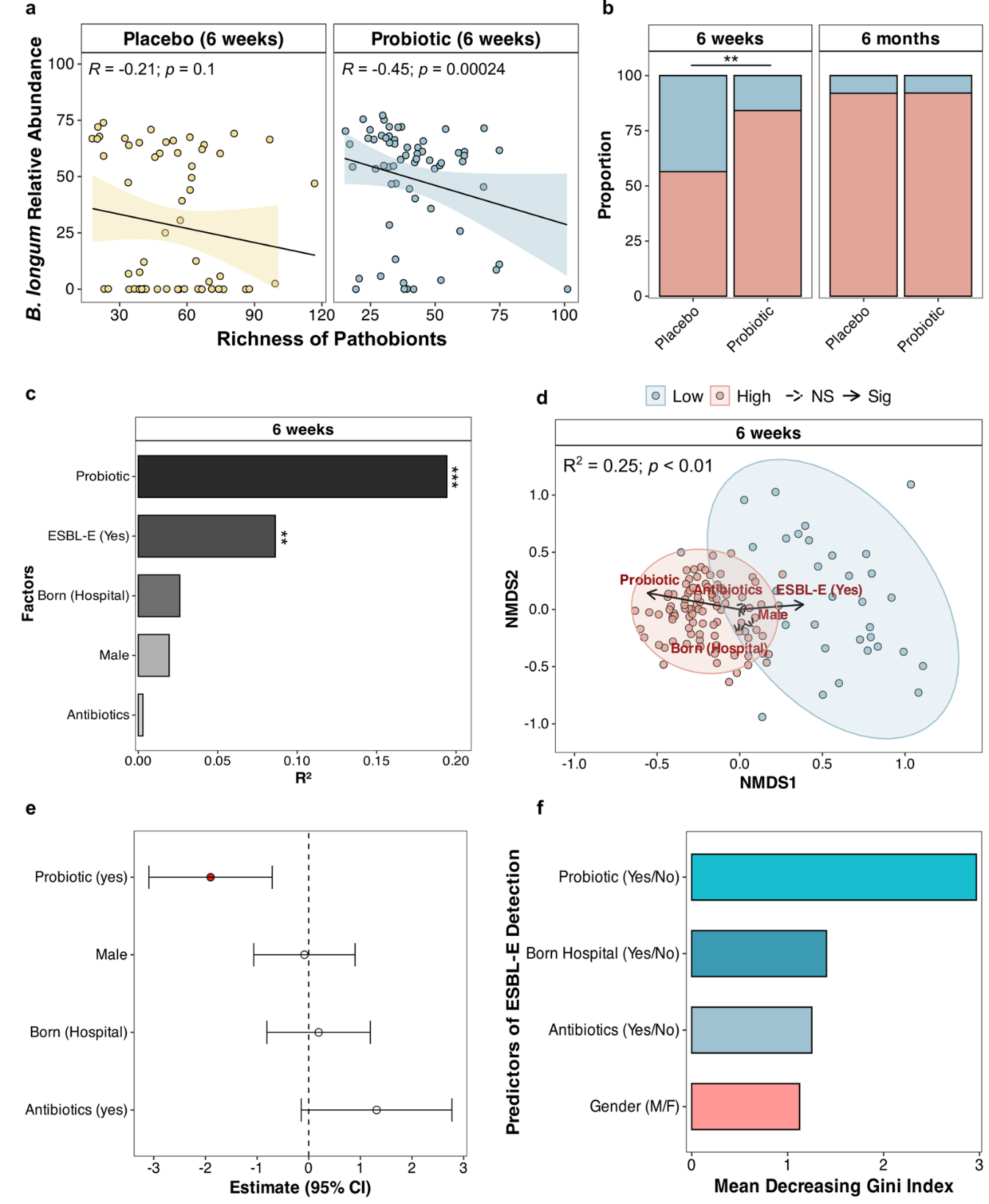

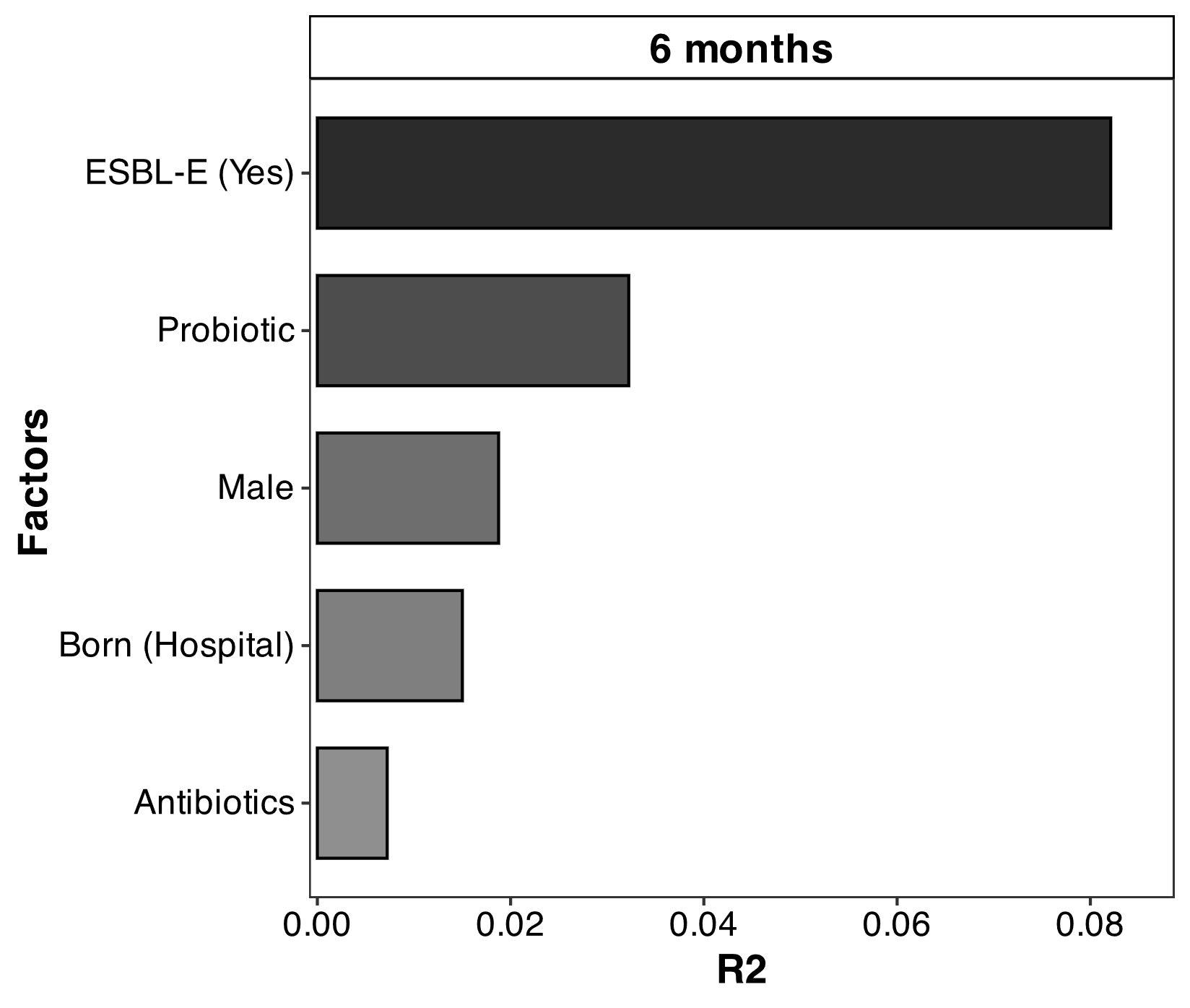

**

**d**

**c**

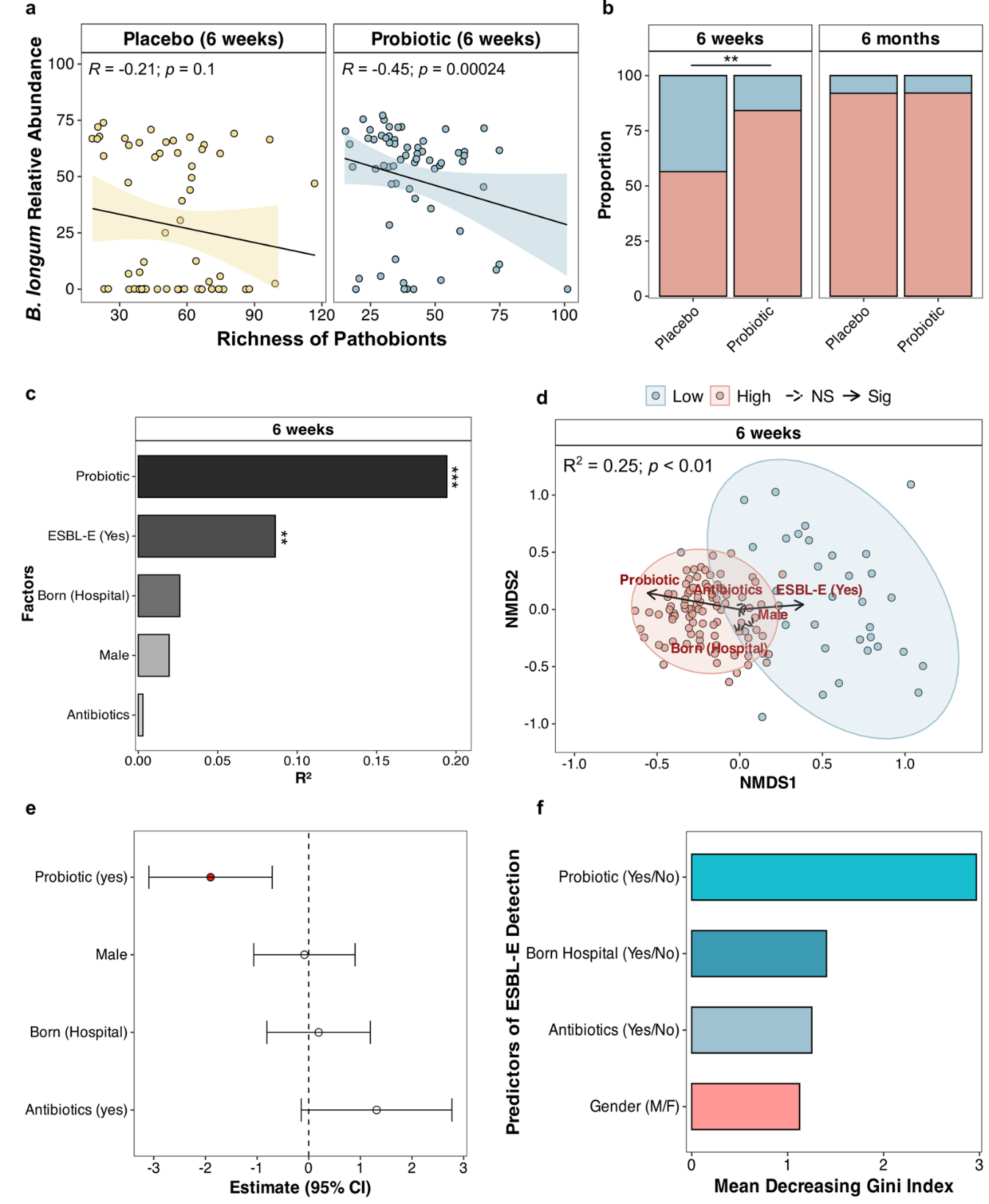
**Figure S3**. **Impact of probiotics on ESBL-E carriage.** The impact of selected variables on the ESBL-E carriage as determined by generalised linear modelling (a) and random forest (b). The random forest analysis does not show which specific binary outcome a variable is correlated with, but it shows which variables are most important for prediction. Statistical significance: *p* < 0.05

**b**

**a**

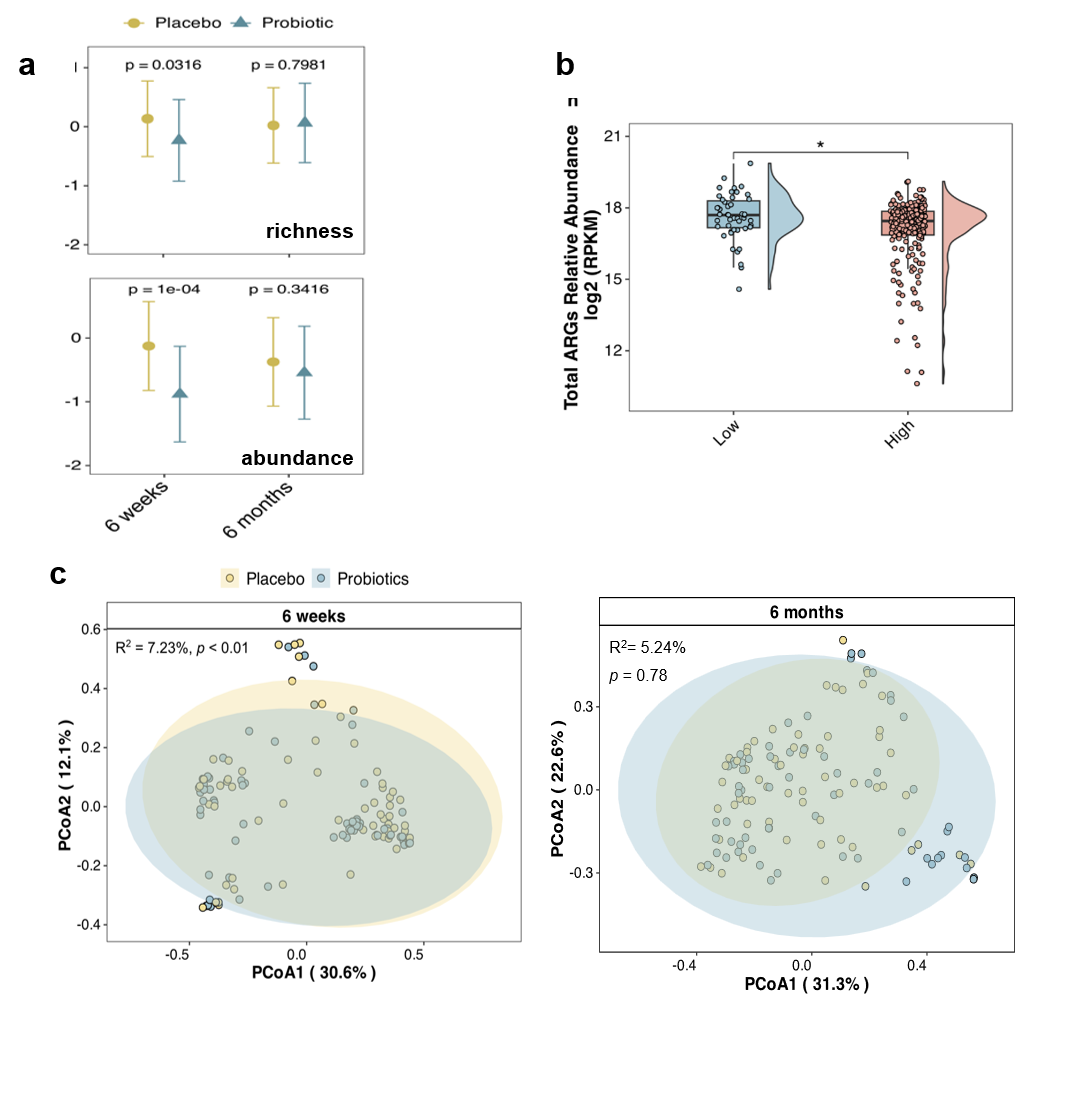
**Figure S4.** **The influence of probiotics on the resistome.** a) Estimated marginal means of probiotic impact on resistome richness and relative abundance across time points. b) Comparison of ARGs relative abundance between microbiota clusters, with each point representing a sample. P-values were computed using the Mann-Whitney U test. c) Principal Coordinate Analysis (PCoA) illustrating resistome beta diversity differences between the probiotic and placebo groups at 6 weeks and 6 months, visualised through Jaccard dissimilarity comparisons. Statistical significance: *p* < 0.05. Symbols: ∗ p < 0.05

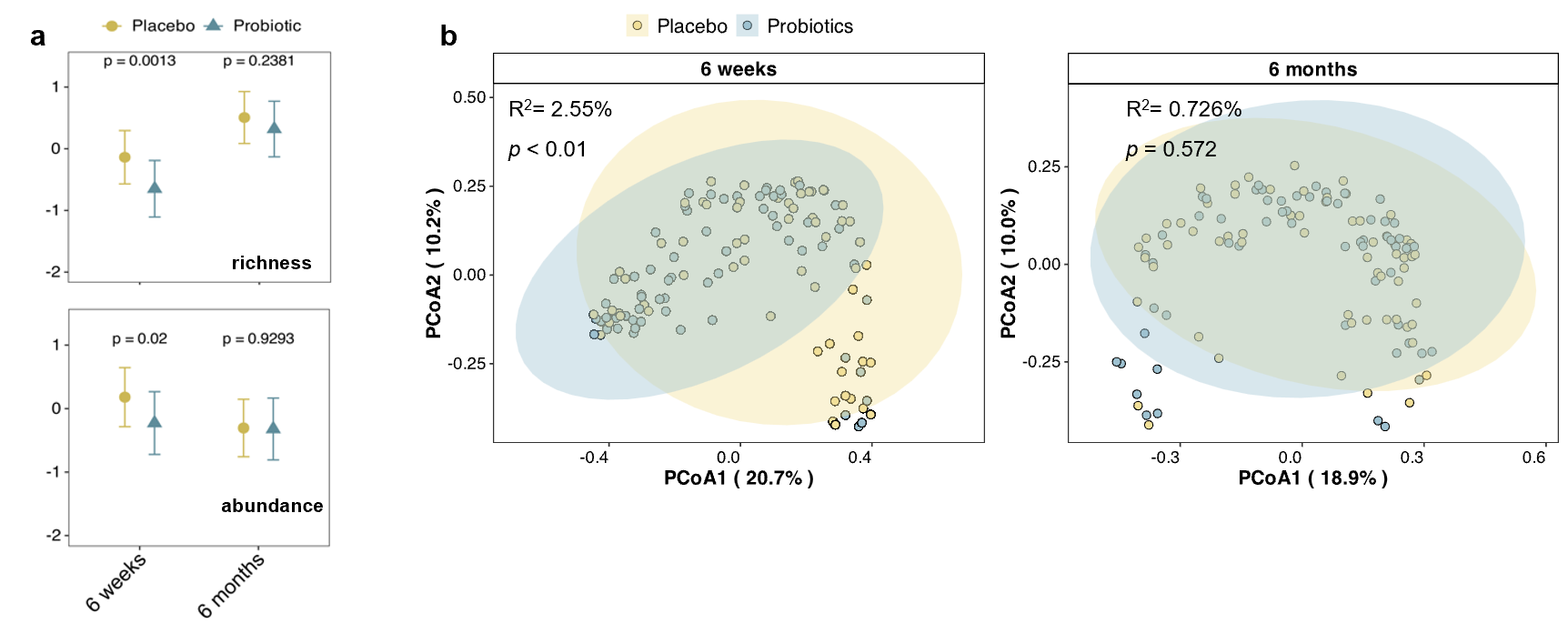

**Figure S5.** **The influence of probiotics on the mobilome.** a) Estimated marginal means of probiotic impact on mobilome richness and relative abundance across time points. b) Principal Coordinate Analysis (PCoA) illustrating mobilome beta diversity differences between the probiotic and placebo groups at 6 weeks and 6 months, visualised through Jaccard dissimilarity comparisons. Statistical significance: *p* < 0.05

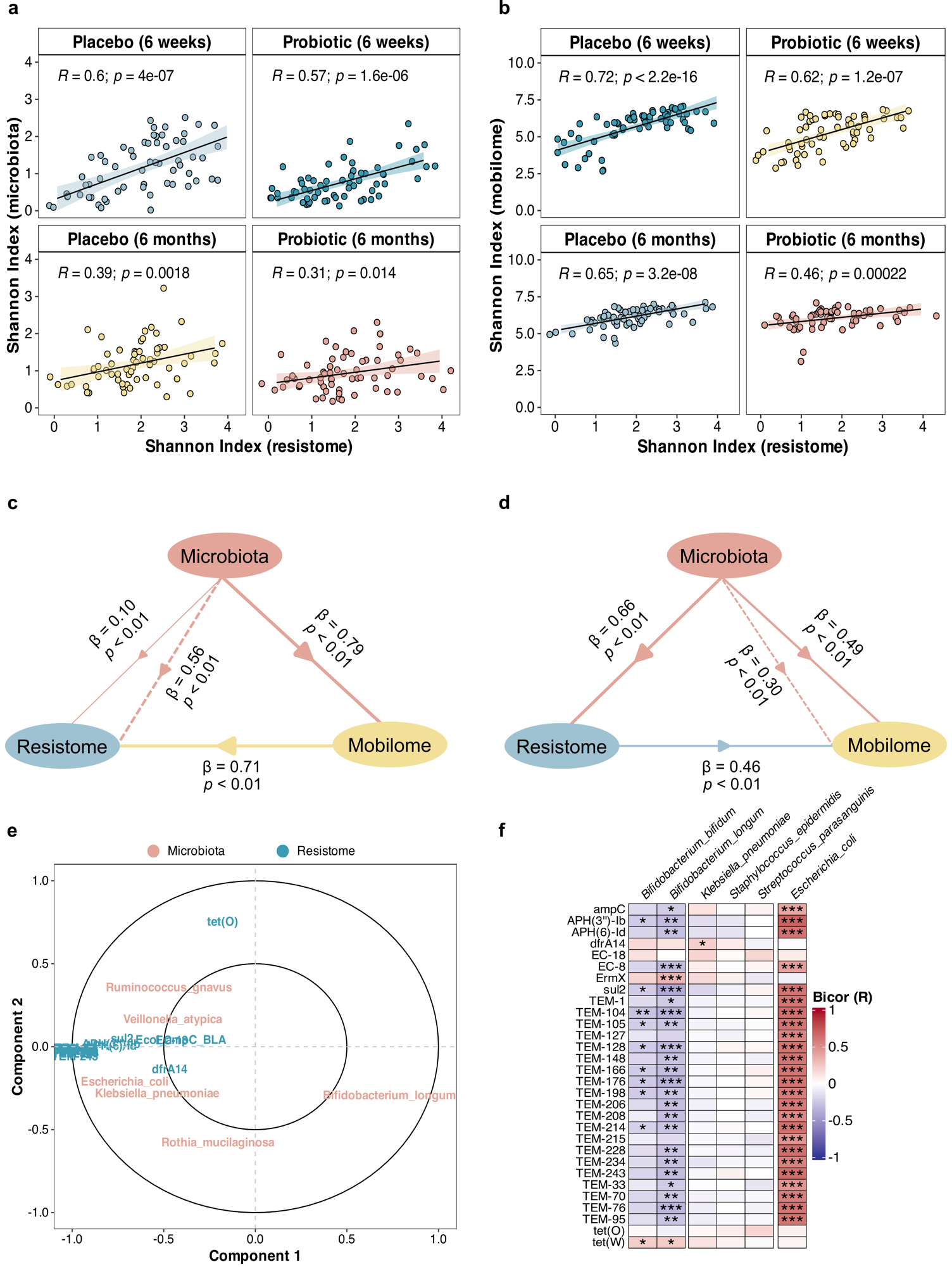
**Figure S6. Correlation analysis.** Spearman correlation analysis between resistome diversity and microbiota diversity (a) and between resistome diversity and mobilome diversity (b). The strength of correlation estimated by R values: 0.10-0.30 weak, 0.30-0.50 moderate, 0.5-0.7 strong, 0.7-1 very strong.

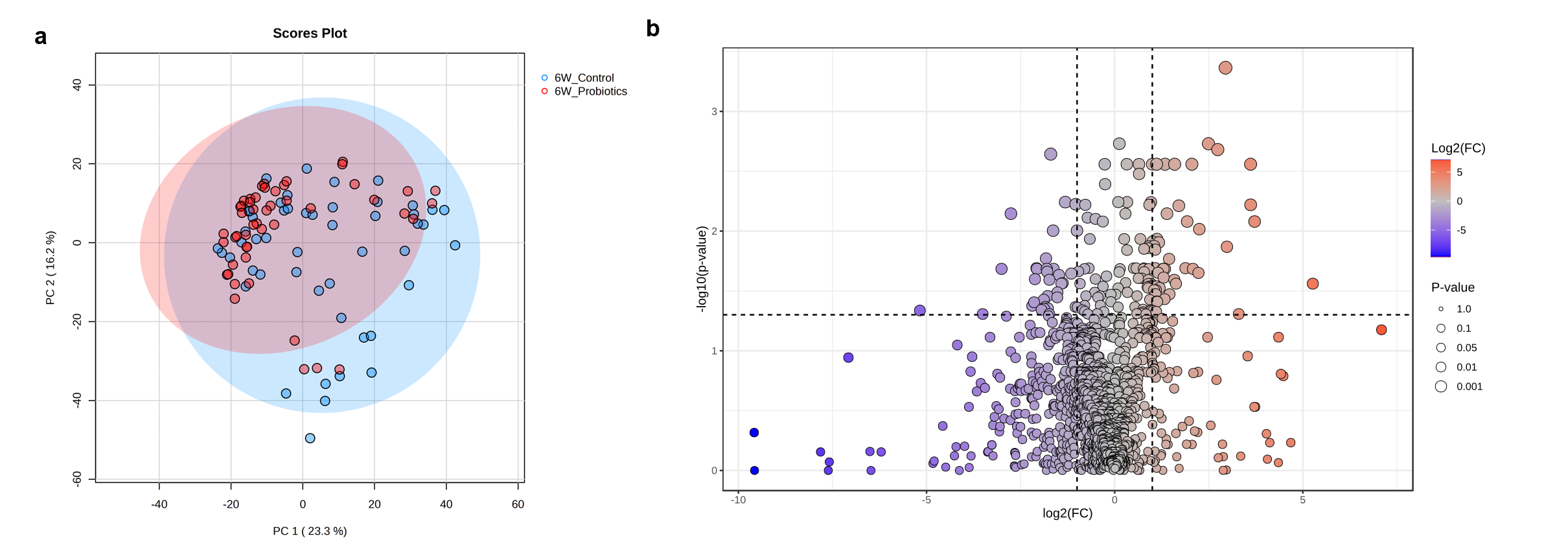
 **Figure S7. Comparison of the fecal metabolome of placebo and probiotic groups at 6 weeks.** A) Principal component analysis (PCA) of 1188 detected metabolic features (R = 0.07218, *p* = 0.004) using Euclidean distance matrix. B) Volcano plot describing differences in metabolic features between probiotic and placebo groups. In the probiotic group, there were 45 features with significantly increased abundance, while 37 features had significantly decreased abundance. Settings used in metaboAnalyst: *p*-value set to 0.05 (Mann-Whitney U test), accounting for unequal group variance, applying false discovery rate correction, and a fold change cutoff of 2.0. For both analyses, the untargeted metabolomic data were normalized by median, log_10_ transformed, and Pareto scaled.

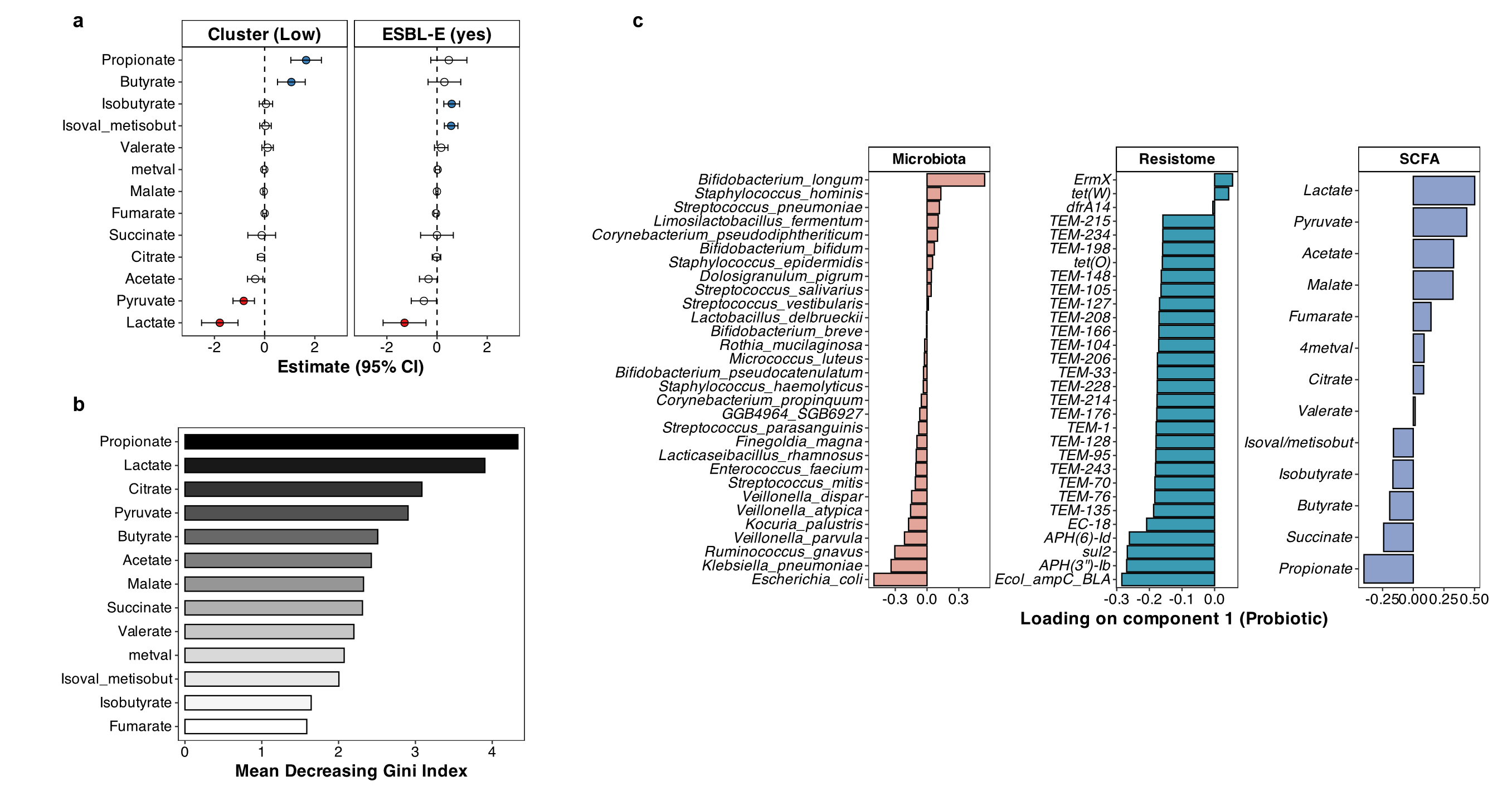
 **Figure S8 Targeted metabolomics data analysis. (a)** Differential abundance of the metabolites identified using linear model (Benjamini-Hochberg adjusted p values). (**b**) The importance of SCFAs on microbiota composition as determined by random forest. (**c**) SCFA loadings from DIABLO analysis for the probiotic component.

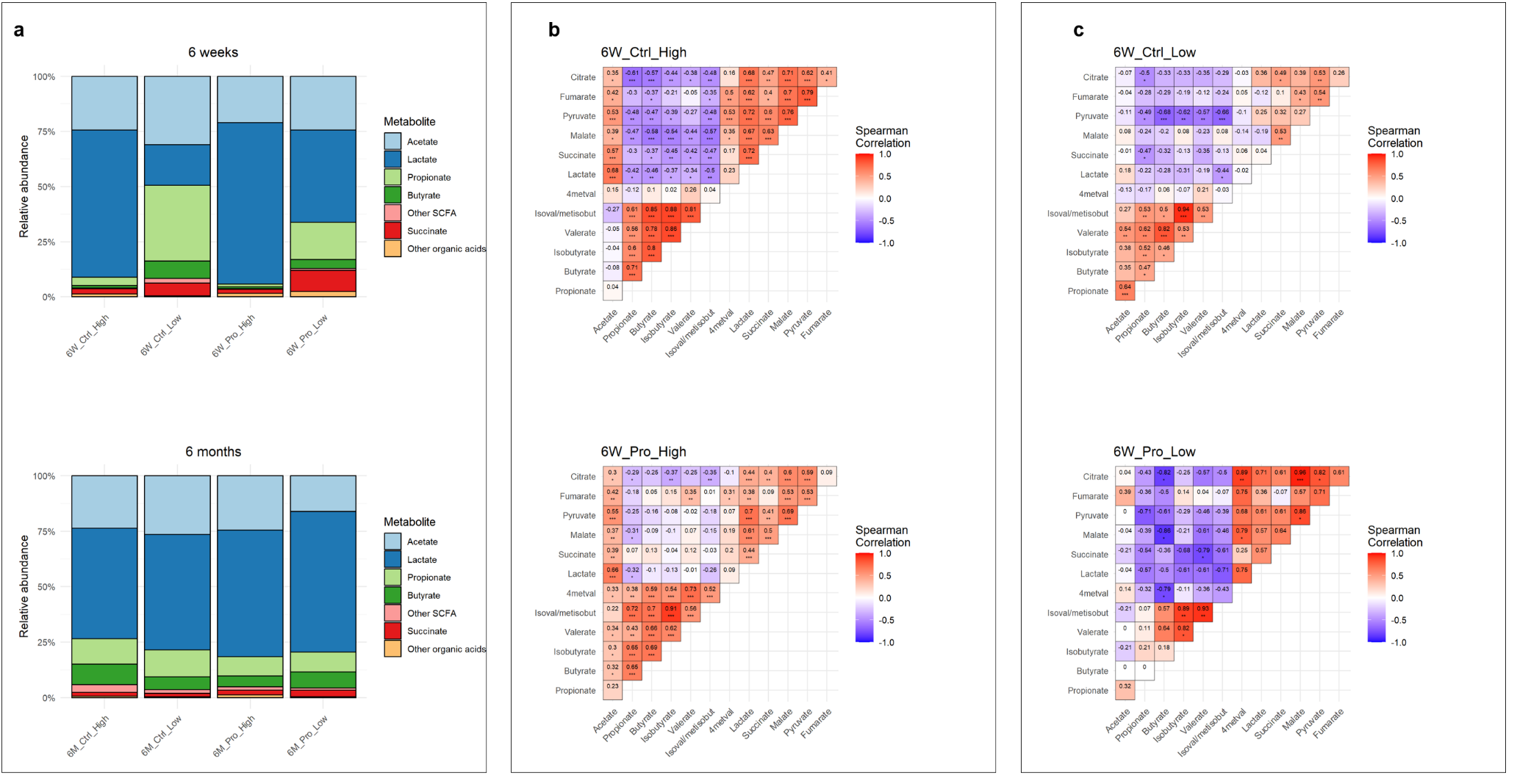
**Figure S9.** **Dynamics of SCFAs and OAs based on *Bifidobacterium* spp. abundance.** a) Relative proportion of SCFAs and organic acids in samples from 6 weeks and 6 months for the probiotic and control group, and with either high or low *Bifidobacterium* spp. abundance. Correlation heatmaps showing Spearman correlations between metabolites for b) "High" and c) "Low" groups at 6 weeks. Asterisks indicate significance levels (*p < 0.05, **p < 0.01, ***p < 0.001).

**Table S1**. Baseline characteristics of participants

| **Variables** | **Probiotics (N=80)** | **Placebo (N=72)** |
| --- | --- | --- |
| **Mother and delivery** |  |  |
| Maternal age, median (IQR), years | 26.5 (22.0 – 31.0) | 25 (22.0 - 29.3) |
| Maternal HIV infection | 5.0 % (4/80) | 2.8 % (2/72) |
| Mode of delivery – n (%) |  |  |
| - Vaginal | 97.5 % (78/80) | 95.8 % (69/72) |
| - Caesarean section | 2.5 % (2/80) | 4.2 % (3/72) |
| Birthplace – n (%) |  |  |
| - Haydom Lutheran Hospital | 40.0 % (32/80) | 43.1 % (31/72) |
| - Other health facility | 25.0 % (20/80) | 22.2 % (16/72) |
| - At home | 35.0 % (28/80) | 34.7% (25/72) |
| **Child** |  |  |
| Female sex – n (%) | 48.8 % (39/80) | 50.0 % (36/72) |
| Birth weight, median (IQR) - kg | 3.3 (3.0 – 3.5) | 3.3 (3.0 – 3.5) |
| Birth length, median (IQR) - cm | 48.0 (47.0 – 49.0) | 48.5 (47.0 – 49.0) |
| Apgar score at 5 min, median (IQR)* | 10 (10-10) | 10 (10-10) |
| Apgar score at 5 min < 9 – n (%)* | 0 (0%) | 0 (0%) |
| Apgar score at 5 min |  |  |
| - 9 | 2 (2.5 %) | 3 (4.2 %) |
| - 10 | 29 (36.3 %) | 27 (37.5 %) |
| - Not recorded | 49 (61.3 %) | 42 (58.3 %) |
| **Breastfeeding** |  |  |
| At 6 weeks – no. (%) † | 100.0 % (80/80) | 100.0 % (72/72) |
| At 6 months – no. (%) † | 95.0 % (76/80) | 94.4 % (68/72) |
| **Caregiver-reported adherence during the 4-week intervention** |  |  |
| Study drug given daily – no. (%) | 95.0 % (76/80) | 84.7 % (61/72) |
| Forgot a dose 1-6 days – no. (%) | 3.8 % (3/80) | 13.9 % (10/72) |
| Forgot a dose 7-21 days – no. (%) | 1.3% (1/80) | 0.0 % (0/72) |
| Forgot a dose ≥ 22 days– no. (%) | 0.0 % (0/80) | 1.4% (1/72) |

| Pathobiont | Source |
| --- | --- |
| *Acinetobacter baumannii* | Shao.et,al (2019); Murray et al., 2022 |
| *Campylobacter jejuni* | Burnham et al, 2018 |
| *Clostridium perfringens* | Shao.et,al (2019) |
| *Enterobacter cloacae* | Shao.et,al (2019) |
| *Enterococcus faecalis* | Shao.et,al (2019); Murray et al., 2022 |
| *Enterococcus faecium* | Shao.et,al (2019) |
| *Escherichia coli* | Murray et al., 2022 |
| *Haemophilus influenzae* | Murray et al., 2022 |
| *Klebsiella oxytoca* | Shao.et,al (2019) |
| *Klebsiella pneumoniae* | Shao.et,al (2019); Murray et al., 2022 |
| *Morganella morganii* | Murray et al., 2022 |
| *Pseudomonas aeruginosa* | Shao.et,al (2019); Murray et al., 2022 |
| *Salmonella enterica* | Murray et al., 2022 |
| *Serratia marcescens* | Murray et al., 2022 |
| *Staphylococcus aureus* | Murray et al., 2022 |
| *Staphylococcus epidermidis* | Shao.et,al (2019) |
| *Streptococcus parasanguinis* | Shao.et,al (2019) |
| *Streptococcus pneumoniae* | Murray et al., 2022 |

**Table S10**. Bacterial species with pathogenic potential reported by previous studies
